# Development of an engineered extracellular vesicles-based vaccine platform for combined delivery of mRNA and protein to induce functional immunity

**DOI:** 10.1101/2024.03.14.585062

**Authors:** Xin Luo, Kathleen M. McAndrews, Kent A. Arian, Sami J. Morse, Viktoria Boeker, Shreyasee V. Kumbhar, Yingying Hu, Krishnan K. Mahadevan, Kaira A. Church, Sriram Chitta, Nicolas T. Ryujin, Janine Hensel, Jianli Dai, Dara P. Dowlatshahi, Hikaru Sugimoto, Michelle L. Kirtley, Valerie S. LeBleu, Shabnam Shalapour, Joe H. Simmons, Raghu Kalluri

**Affiliations:** Department of Cancer Biology, University of Texas MD Anderson Cancer Center, Houston, TX; Department of Bioengineering, Rice University, Houston, TX; Michale E. Keeling Center for Comparative Medicine and Research, University of Texas MD Anderson Cancer Center, Bastrop, TX; Department of Medicine, Baylor College of Medicine, Houston, TX; Department of Molecular and Cellular Biology, Baylor College of Medicine, Houston, TX

## Abstract

mRNA incorporated in lipid nanoparticles (LNPs) became a new class of vaccine modality for induction of immunity against COVID-19 and ushered in a new era in vaccine development. Here, we report a novel, easy-to-execute, and cost effective engineered extracellular vesicles (EVs)-based combined mRNA and protein vaccine platform (EV^X-M+P^ vaccine) and explore its utility in proof-of-concept immunity studies in the settings of cancer and infectious disease. As a first example, we engineered EVs to contain ovalbumin mRNA and protein (EV^OvaM+P^) to serve as cancer vaccine against ovalbumin-expressing melanoma tumors. EV^OvaM+P^ administration to mice with established melanoma tumors resulted in tumor regression associated with effective humoral and adaptive immune responses. As a second example, we generated engineered EVs, natural nanoparticle carriers shed by all cells, that contain mRNA and protein Spike (S) protein to serve as a combined mRNA and protein vaccine (EV^SpikeM+P^ vaccine) against SARS-CoV-2 infection. EV^SpikeM+P^ vaccine administration in mice and baboons elicited robust production of neutralizing IgG antibodies against RBD (receptor binding domain) of S protein and S protein specific T cell responses. Our proof-of-concept study describes a new platform with an ability for rapid development of combination mRNA and protein vaccines employing EVs for deployment against cancer and other diseases.

## Introduction

Extracellular vesicles (EVs) are small vesicles secreted by presumably all types of cells (Kalluri and LeBleu 2020). EVs consist of two major subtypes, exosomes and ectosomes, and contain a lipid bilayer resulting from their origin from either the endosomal pathway or direct budding from the plasma membrane (Kalluri and McAndrews 2023). Exosomes are presumably derived from the endosomal pathway and have a size range of approximately 40 to 160 nm in diameter. Ectosomes are generated from the direct outward budding of the cell membrane with a size range of approximately 50 nm to 1 µm in diameter (Kalluri and LeBleu 2020). The precise physiological role of EVs remains unknown, but they are postulated to function in cargo transfer between cells, such as DNA, RNA, lipids, proteins, and metabolites, and participate in intracellular communication (Yáñez-Mó, Siljander et al. 2015, Van Niel, d’Angelo and Raposo 2018, Kalluri and McAndrews 2023). The biocompatible nature of EVs imparts advantages for in vivo delivery of therapeutic payloads for the treatment of different diseases (Lener, Gimona et al. 2015). Dendritic cells derived EVs loaded with cancer antigen peptides have been used to treat melanoma and small cell lung cancer and demonstrated inhibition of the tumor growth (Escudier, Dorval et al. 2005, Morse, Garst et al. 2005). EVs have also been reported to deliver small interfering RNAs and microRNAs targeting cancer oncogenes with great success (Ohno, Takanashi et al. 2013, Kamerkar, LeBleu et al. 2017, Mendt, Kamerkar et al. 2018, Haltom, Hassen et al. 2022).

Based on the broad applications and successful implementation of EVs as therapies in different disease contexts, we aimed to generate an easy to execute EV-based vaccine platform where antigens that drive disease can be loaded into EVs and conducted proof-of-concept studies. To establish the efficacy of such EV-based vaccine platform (EV^X-M+P^), we evaluated immune responses to EV vaccination embedded with two different antigens, the model antigen ovalbumin (OVA) and the S protein of SARS-CoV-2. OVA has been widely employed in the context of cancer immunotherapy as a model antigen (Xu, Lv et al. 2020), especially when combined with adjuvant or in nanovaccine delivery (Liu, Xie and Zheng 2022). The isolation, structure, and amino acid sequence of OVA was established many years ago, enabling its use in early studies of antibody generation (Hopkins 1900, Heidelberger, Pedersen and Tiselius 1936, Huntington and Stein 2001). Moreover, the mechanisms of OVA antigen presentation, OVA peptide epitopes, and humoral and cellular immune responses are well-characterized (Beck and Spiegelberg 1989, Rotzschke, Falk et al. 1991, Ke, Li and Kapp 1995, Dudani, Chapdelaine et al. 2002). As a result, we employed EV-associated OVA as a model vaccination strategy to evaluate its feasibility and efficacy in cancer prevention. We hypothesized that EVs could be engineered to generate antigen-specific immune responses with diverse antigens.

Severe Acute Respiratory Syndrome Coronavirus 2 (SARS-CoV-2) was identified as the virus responsible for the COVID-19 pandemic (Wu, Leung and Leung 2020). S protein is one of the four structure proteins of SARS-CoV-2 (Harrison, Lin and Wang 2020, Naqvi, Fatima et al. 2020) and allows for docking of the virus with the cell surface via binding to the cell surface receptor angiotensin converting enzyme 2 (ACE2) through its receptor binding domain (RBD) to release the genome of the virus to the host cytosol, making it an important component for host specificity (Wang, Zhang et al. 2020). To date, EVs have been investigated for the therapeutic potential against SARS-CoV-2, with a focus on utilizing ACE2 carrying EVs as nanodecoys to neutralize the virus (Li, Wang et al. 2021, El-Shennawy, Hoffmann et al. 2022). S protein has been utilized as a target for antibody production and vaccine development due to its host specificity and the potential to generate neutralizing antibodies against S protein to prevent viral entry into cells (McAndrews, Dowlatshahi et al. 2020). However, wildtype S protein has been reported to be unstable. A two proline (2P) amino acid substitution in the site of 986 and 987 between heptad repeat 2 (HR2) and central helix (CR) of S protein stabilizes the metastable S protein and increases its half-life by preventing the cleavage between S1 and S2 subunits (Pallesen, Wang et al. 2017, Corbett, Edwards et al. 2020, Lee, Kim et al. 2021). Moreover, the prefusion-stabilized S protein is known to be more immunogenic than the wildtype form (Pallesen, Wang et al. 2017, Corbett, Edwards et al. 2020).

In this study, we developed EV-based vaccines platform (EV^X-M+P^) where X can be any antigen/s related to infectious diseases or cancers and M+P representing a combination of antigenic protein for direct antigen presentation and mRNA which could be delivered to target cells and potentially translated into additional antigenic protein. Specifically, the ability of such EV-based platform to generate potentially protective humoral and cellular adaptive immune responses was demonstrated in two contexts: 1) EV^OvaM+P^ delivering OVA and 2) EV^SpikeM+P^ delivering S protein modified with a 2-proline substitution (S(2P)-protein).

## Results

### The generation of EV^OvaM+P^ and efficacy as a therapeutic vaccine against melanoma tumors

In order to evaluate EVs as vaccine candidates, 293F cells, an FDA approved cell line for the generation of a number of biologics, were stably transduced to overexpress OVA. While it is an attractive option to use protein or lipid anchors to display antigens using complex engineering techniques and employ microfluid devices to isotate EVs for translational applications, it comes with a high cost and minimal opportunity for easy adaptability and common use (Cheng and Kalluri 2023, Kalluri and McAndrews 2023). We used simple plasmid transfection to express OVA or SARS-CoV2 S protein in 293F and isolated EVs that contained the given mRNA and protein and evaluated them as vaccine candidates. Such engineered EVs isolated from 293F-OVA cells were termed EV^OvaM+P^. Nanoparticle tracking analysis (NTA) was performed to show the size distribution and concentration of the vesicles (**Figure 1a**). With the average particle diameters measuring 134 nm and 133 nm for control EVs and EV^OvaM+P^, respectively, the presence of OVA did not alter the size of the vesicles (**Figure 1a**). Moreover, OVA did not impact EVs production, as measured by NTA and EV protein content (**Figure 1b-c**). To characterize the biological properties of EV^OvaM+P^, flow cytometry was performed to validate the putative exosomes biomarkers (CD9, CD47, CD63, CD81) (**Figure 1d**). Our results also indicate that EVs produced from 293F cells present with CD47, the ‘don’t eat me’ signal previously described on EVs used in therapeutic payload delivery that inhibits EV recognition and clearance by monocytes in circulation (Kamerkar, LeBleu et al. 2017, Mendt, Kamerkar et al. 2018). Collectively, these data confirm that EV^OvaM+P^ have similar biological properties as unmodified EVs. The presence of OVA mRNA in EV^OvaM+P^ was validated by qRT-PCR (**Figure 1e**) and OVA protein detected by western blot in the EV-producing 293F cells (**Figure 1f**, **Figure S1a**) and EV^OvaM+P^ (**Figure 1g**, **Figure S1b**). Approximately 3% of total EV^OvaM+P^ protein consists of OVA (**Figure S1b**).

**Figure 1.**
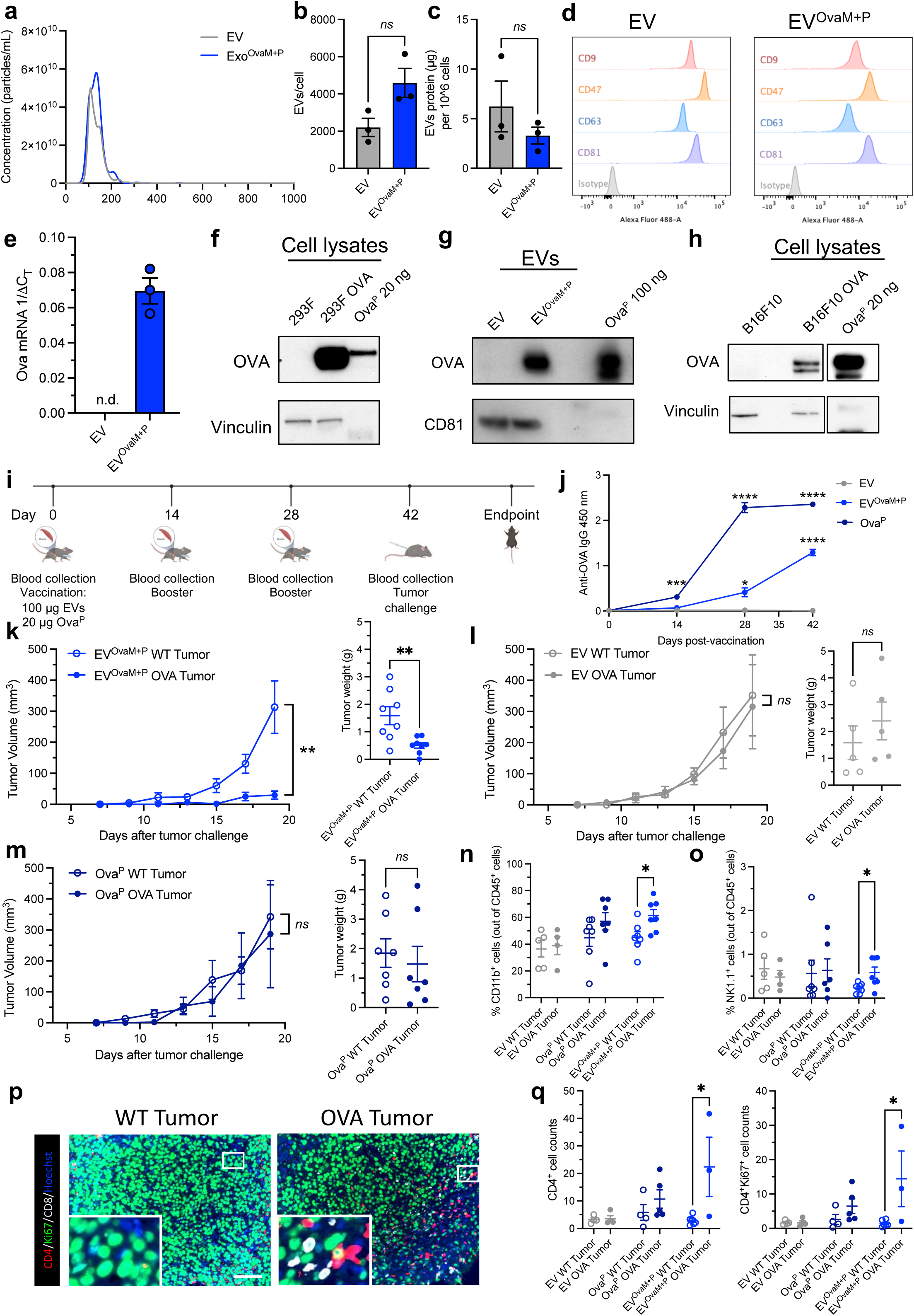
EV^OvaM+P^ elicits antigen specific immune responses to prevent tumor growth. (**a**) NTA of wide type 293F EV and EV^OvaM+P^. (**b-c**) EVs production from cells quantified by NTA (**b**) and microBCA assay(**c**) (n = 3). (**d**) Flow cytometry analysis of EVs markers CD9, CD47, CD63, and CD81 (n = 3). (**e**) Quantification of OVA mRNA from EV^OvaM+P^. Data were shown as 1/-1CT (n = 3). n.d., not detected. (**f-g**) Representative western blot of OVA protein expression in 293F OVA^+^ cells and EV^OvaM+P^ (n = 3). (**h**) Representative western blot of OVA protein expression in B16F10 OVA cells (n = 3). (**i**) Schematic illustration of vaccination and tumor challenge schedule with EV^OvaM+P^ in mice. C57BL/6J mice were intramuscularly injected with EVs, EV-SC2S and EV^OvaM+P^ and recombinant OVA protein (OVA^P^). (**j**) Quantification of anti-OVA IgG antibodies from vaccinated C57BL/6J mice by ELISA (n = 6 per group). (**k**) Tumor volume and weight measurement of wide type and OVA tumor in EV^OvaM+P^ vaccinated mice (n = 6 per group). (**l**) Tumor volume and weight measurement of wide type and OVA tumor in EV vaccinated mice (n = 8 per group). (**m**) Tumor volume and weight measurement of wide type and OVA tumor in OVA^P^ vaccinated mice (n = 7 per group). (**n**) Frequency of CD11b^+^ cells out of CD45^+^ cells from tumor tissue (n = 6 for EV vaccination, n = 8 for EV^OvaM+P^, n = 7 for OVA^P^). (**o**) Frequency of NK1.1^+^ cells out of CD45^+^ cells from tumor tissue (n = 6 for EV vaccination, n = 8 for EV^OvaM+P^, n = 7 for OVA^P^). (**p**) Representative images of CD4^+^ T cell and CD4^+^ T cell proliferation by TSA staining in WT tumor and OVA tumor tissue from EV^OvaM+P^ vaccinated mice. Scale bar, 100 μm. (**q**) Quantification of CD4^+^ T and proliferative CD4^+^ T cells (Ki67^+^) of (**p**). * P<0.05, ** P<0.01, *** P<0.001, **** P<0.0001 ns: not significant.

With the fabrication of EV^OvaM+P^, we next evaluated EV^OvaM+P^ as a vaccine and its ability to generate effective immune responses. C57BL/6J mice were vaccinated with alhydrogel adjuvant with 100 μg of EV, 100 μg of EV^OvaM+P^ (approximately 3 μg of OVA protein, quantified by western blot (**Figure S1b**)), or 20 μg of recombinant OVA protein (OVA^P^), respectively (details described in the methods, schematic illustration shown in **Figure 1h**). After vaccination, mice were challenged with B16 wildtype (WT) tumor in one flank and B16-OVA tumor in the other flank. Expression of OVA in B16-OVA cells was validated by western blot (**Figure 1i**). IgG antibodies against OVA were detected by ELISA after the first vaccination of OVA^P^ (Day 14), increased after the first booster of EV^OvaM+P^ (Day 28), and maintained until the tumor challenge (Day 42, **Figure 1j**). The growth of the OVA tumor was significantly delayed in the EV^OvaM+P^ vaccinated mice compared to the WT tumor (**Figure 1k**), while in the EV vaccinated (**Figure 1l**) and OVA^P^ vaccinated (**Figure 1m**) mice, insignificant differences between the growth of WT and OVA tumors were observed. Although OVA specific IgG antibodies were detected in the OVA^P^ vaccinated mice (**Figure 1j**), indicating induction of humoral immunity, it was not sufficient to inhibit tumor growth (**Figure 1m**).

Next, we evaluated cellular responses to EV vaccination. The presence of myeloid cells (CD11b^+^) and NK cells (NK1.1^+^) were significantly increased in the OVA tumors of EV^OvaM+P^ vaccinated mice compared to the corresponding WT tumor (**Figure 1n-o**). In contrast, EV or Ova^P^ vaccination did not impact the relative proportion of CD11b^+^ or NK1.1+ cells in OVA tumors (**Figure 1n-o**). CD11b^+^ cells can act as antigen presenting cells, suggesting that the increased abundance of myeloid cells may result in antigen presentation and activation of T cells. As a result, we evaluated the presence of CD4^+^ and CD8^+^ T cells and their activation/proliferation status with Ki67. Immunostaining showed significantly increased proliferative CD4^+^ T cells (Ki67^+^) in the TME of the OVA compared to WT tumor of EV^OvaM+P^ vaccinated mice (**Figure 1p-q, Figure S2**), suggesting an activated T cell response specific to EV^OvaM+P^ vaccination. All these results together indicate that EV^X-M+P^ is able to elicit humoral and cellular responses to suppress tumor growth.

### Generation and validation of EV, EV-SC2S, and EV^SpikeM+P^

We next evaluated the EV^X-M+P^ platform in the context of infectious diseases. To generate EV-SC2S and EV^SpikeM+P^, 293F cells were stably transfected to express S protein or the prefusion-stabilized S(2P) protein. Engineered EVs isolated from these cells were termed EV-SC2S (overexpressing S protein) and EV^SpikeM+P^ (overexpressing S(2P) protein).. TEM analysis of the engineered EVs demonstrated a characteristic cup-shaped morphology of EVs (**Figure 2a**). The size and concentration were measured by NTA. EV-SC2S and EV^SpikeM+P^ displayed an averaged particle diameter of 143 nm and 140 nm, respectively, characteristic of small EVs and were similar in size to EV (157 nm, **Figure 2a**). EV production (EV per cell) was unchanged in EV-SC2S and EV^SpikeM+P^ when compared with control EV (**Figure 2b**). EV protein assessed by microBCA and normalized to cell numbers showed similar production in EV, EV-SC2S, and EV^SpikeM+P^ producing cells (**Figure 2c**). Expression of putative exosomes biomarkers (CD9, CD47, CD63, CD81) were validated by flow cytometry (**Figure 2d**). qRT-PCR and western blot analysis revealed presence of S protein mRNA and S protein, respectively, in EV-SC2S and EV^SpikeM+P^ (**Figure 2e-f, Figure S4**).

**Figure 2.**
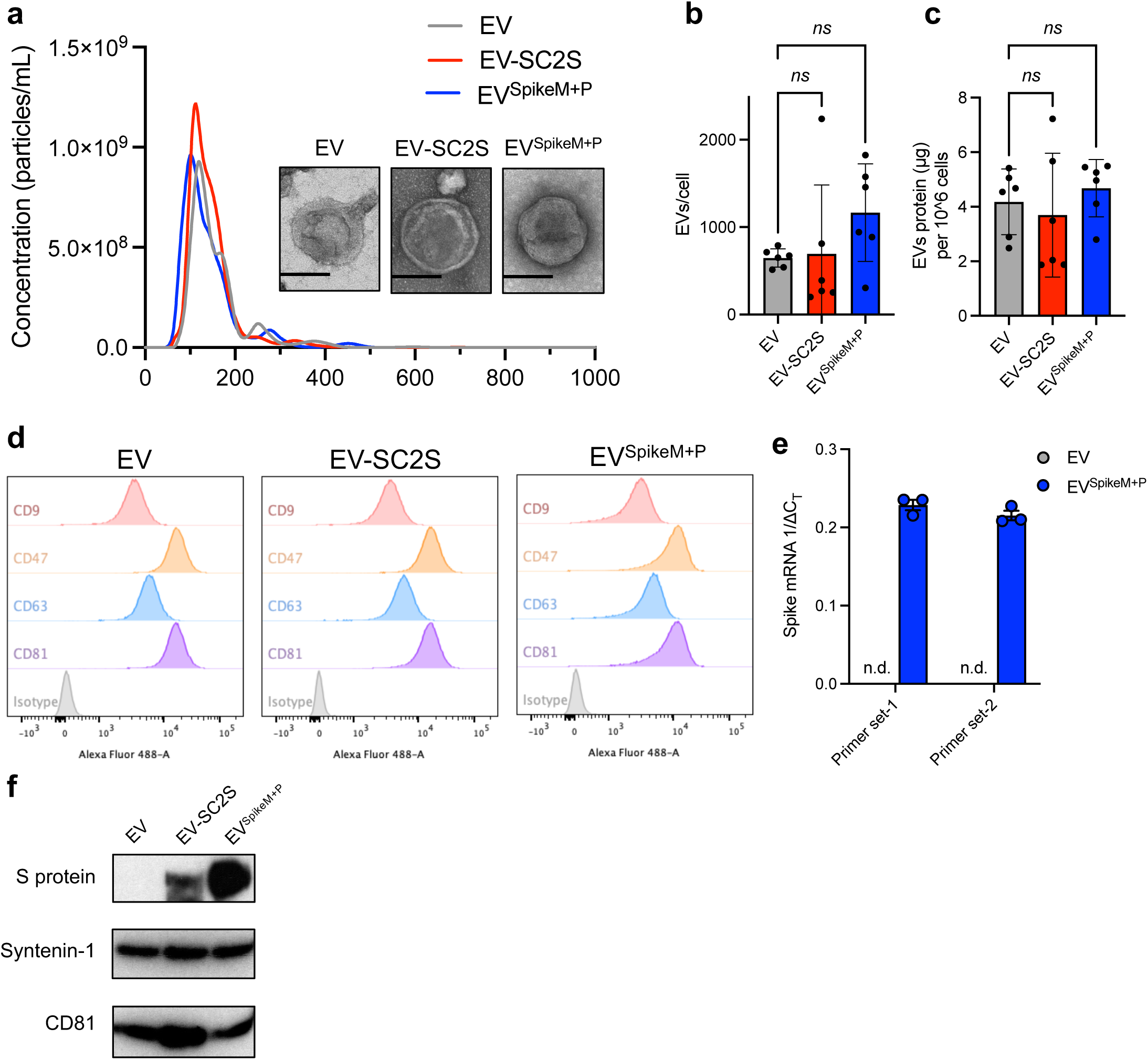
Generation and validation of EV^SpikeM+P^. (**a**) NTA of wild type 293F EV, EV-SC2S, and EV^SpikeM+P^. TEM images of EVs (inset). Scale bar, 100 nm. (**b-c**) EVs production from cells quantified by NTA (**b**) and microBCA assay (**c**) (n = 6). (**d**) Flow cytometry analysis of EVs markers CD9, CD47, CD63, and CD81 (n = 3). (**e**) Quantification of S mRNA from EV^SpikeM+P^. Data were shown as 1/-1CT (n = 3). n.d., not detected. (**f**) Representative western blot of S protein expression in EV-SC2S and EV^SpikeM+P^ (n = 3). ns: not significant.

### SARS-CoV-2 S protein-specific immune response is generated in EV^SpikeM+P^ vaccinated mice

In order to evaluate the efficacy of S(2P) present on EVs in S protein-specific humoral immune response and neutralizing antibodies generation, Balb/c mice were vaccinated with 100 μg of EVs and alhydrogel adjuvant (details described in the methods, **Figure 3a**). No significant changes in body weight were observed during the course of the vaccination and monitoring period in mice administered EV-SC2S or EV^SpikeM+P^ compared to EV (**Figure 3c**). IgG antibodies against RBD were detected at day 28, and with increasing levels over time, specifically in EV^SpikeM+P^ vaccinated mice compared to EV and EV-SC2S vaccinated mice (**Figure 3d**). A neutralizing assay using VSVΔG pseudotyped with S protein on the surface (VSVΔG-Luc-Spike*) was used to evaluate the neutralizing capacity of the antibodies detected in the plasma, where a decreased luminescence indicates pseudovirus neutralization (**Figure 3b**). The antibodies generated by mice in the EV^SpikeM+P^ vaccination group showed significant neutralization of S protein pseudovirus entry into cells (**Figure 3e**).

**Figure 3.**
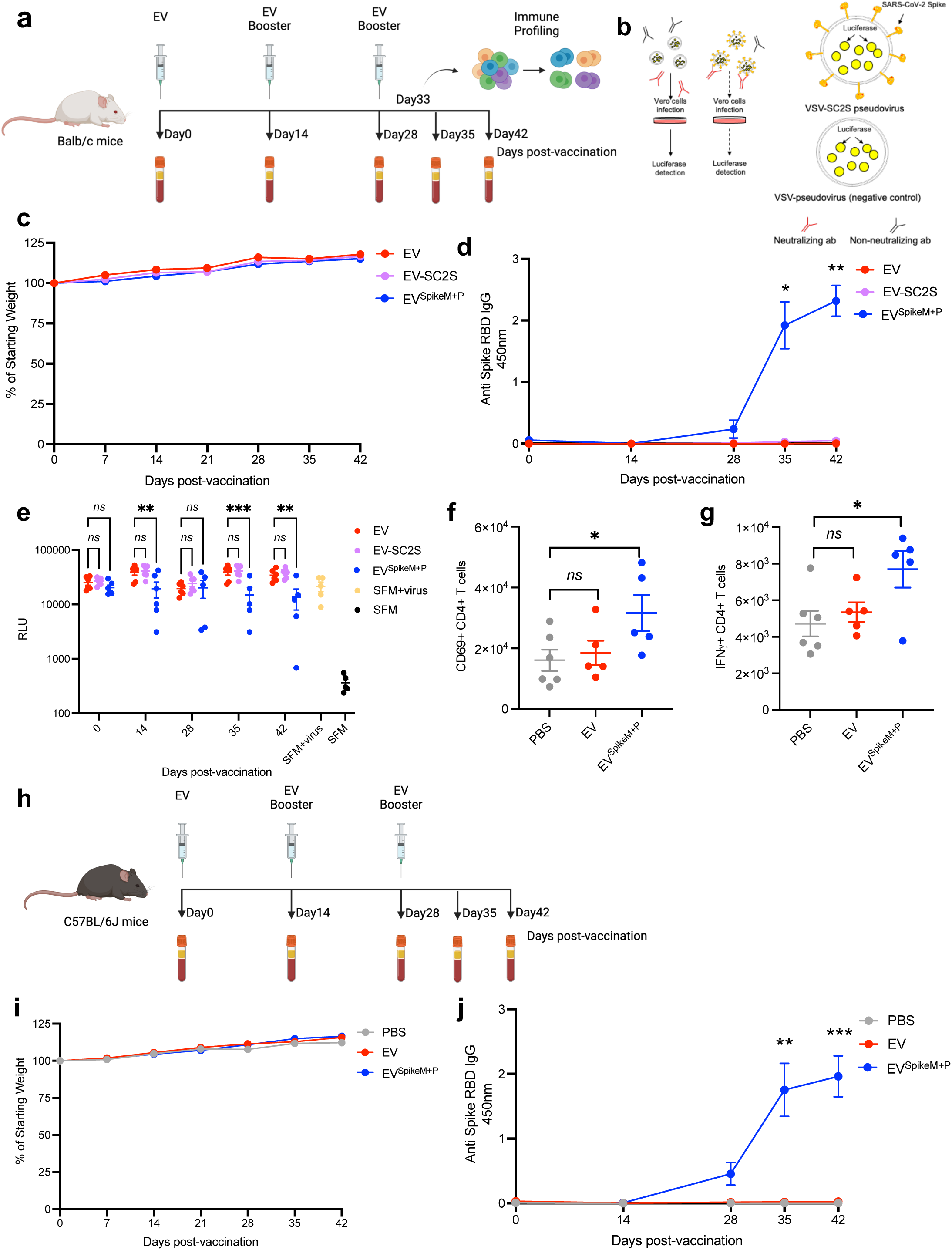
EV^SpikeM+P^ vaccination elicits humoral and cellular responses in mice. (**a**) Schematic illustration of vaccination schedule with EV^SpikeM+P^ in mice. Balb/c micewere intramuscularly injected with EVs, EV-SC2S and EV^SpikeM+P^. (**b**) Schematic illustration of the pseudovirus neutralization assay. (**c**) Body weight change of Balb/c mice during vaccination expressed as percentage of starting weight (n = 5 per group). (**d**) Quantification of anti-S-RBD IgG antibodies from vaccinated Balb/c mice by ELISA (n = 5 per group). (**e**) Neutralizing capacity of the antibodies from vaccinated mice evaluated with pseudovirus neutralization assay (n = 5 per group). Data are expressed as relative luminescence units (RLU). SFM+virus, serum-free media with pseudovirus; SFM, serum-free media alone. (**f-g**) S protein-specific CD4^+^ T cells in Balb/c mice splenocytes showing T cell activation (**f**) and Th1 response (**g**) by intracellular cytokine staining (n = 5 per group). Splenocytes were re-stimulated with S peptide pool. (**h**) Schematic illustration of vaccination schedule with EV^SpikeM+P^ in mice. C57BL/6J mice were intramuscularly injected with PBS, EVs, EV^SpikeM+P^. (**i**) Body weight change of C57BL/6J mice during vaccination expressed as percentage of starting weight (n = 10 per group). (**j**) Quantification of anti-S-RBD IgG antibodies from vaccinated C57BL/6J mice by ELISA (n = 10 per group). * P<0.05, ** P<0.01, *** P<0.001, ns: not significant.

Next, we investigated T cell responses to EV^SpikeM+P^ . Splenocytes were collected 5 days after the last booster (Day 33) (**Figure 3a**) and stimulated with S peptide pool. S protein-specific activation of T cells was observed (**Figure 3f, S4**) and a Th1 immune response was induced by EV^SpikeM+P^ in mice, exemplified by increased IFNγ (**Figure 3g**). We further explored the vaccination efficacy in C57BL/6J mice (**Figure 3h**). Body weight was monitored and similarly, no significant changes were observed in the EV or EV^SpikeM+P^ vaccinated mice compared with the PBS injected mice (**Figure 3i**). Starting from Day 28, S-RBD specific IgG antibodies were detected, specifically in EV^SpikeM+P^ vaccination group (**Figure 3j**). These results indicated the anti-S protein effect induced by the vaccination of EV^SpikeM+P^ in mice, in both Balb/c and C57BL/6J backgrounds.

### EV^SpikeM+P^ elicits S protein-specific immune responses in baboons

Having observed adaptive immune responses in mice after the EV^SpikeM+P^ administration, we wanted to validate the EV^SpikeM+P^ strategy in non-human primates. Olive baboons were vaccinated with EV^SpikeM+P^ at 3 dosages, 0.625 mg, 1.25 mg, 2.5 mg, together with 2% alhydrogel (1:1 ratio) as adjuvant for all 3 immunizations in total administered at a 21 day interval (**Figure 4a**). Blood was collected before and after immunization, to evaluate long-term immune responses (details shown in the schematic **Figure 4a**). IgG antibodies against S-RBD were detected in the higher dosage groups (1.25 mg and 2.5 mg) after the first booster (day 42 post vaccination). The level of IgG antibody was maintained after the second booster (day 79, **Figure 4b**). The titers of the IgG antibodies were measured from the baboons vaccinated with higher dosages of EV^SpikeM+P^ (1.25 mg and 2.5 mg) and decreased around 120 days after the vaccination (**Figure 4c**). We next explored the T cell responses from the vaccinated baboons.

**Figure 4.**
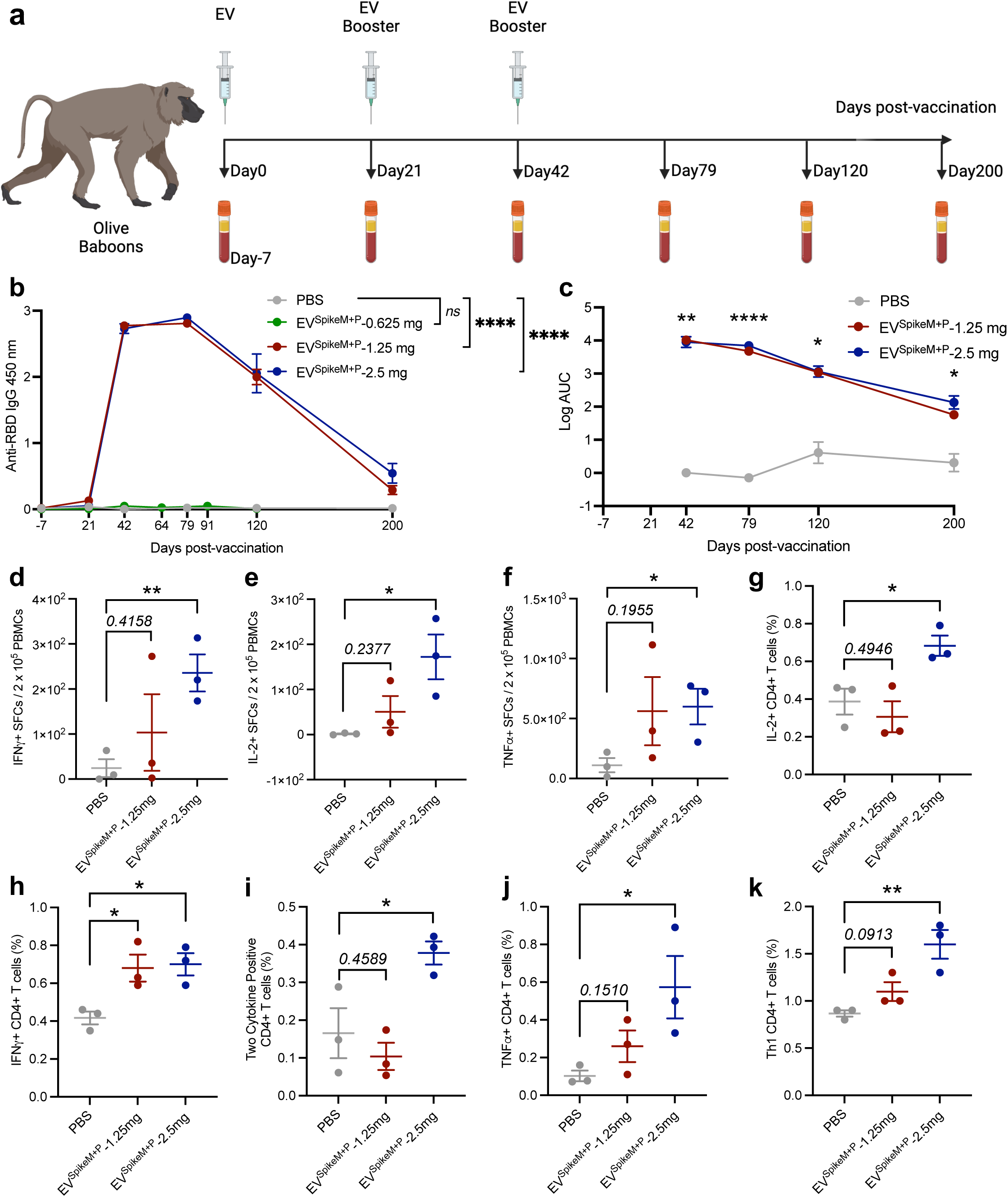
EV^SpikeM+P^ vaccinated baboons develop antibodies and T cell responses against S protein. (**a**) Schematic illustration of vaccination schedule with EV^SpikeM+P^ of 3 dosages in baboons. (**b**) Quantification of anti-S-RBD IgG antibodies from vaccinated baboons by ELISA (n = 3 per group). (**c**) Titration of anti-S-RBD IgG antibodies at different time points plotted as log AUC (area under curve, n = 3 per group). (**d-f**) Quantification of cytokines (IL-2, IFNγ, and TNFα) releasing spots from S protein-specific T cells in circulation re-stimulated with S peptide pool (**d**) or recombinant S protein (**e-f**) by ELISpot (n = 3 per group). Data are shown as SFCs per 2 x 10^6^ cells. (**g-i**) Frequency of S protein-specific CD4^+^ T cells in circulation producing single (IL-2 (**g**), IFNγ (**h**)) and two of Th1 cytokines (IL-2, IFNγ, and TNFα, (**i**)) determined by intracellular cytokine staining (n = 3 per group). PBMCs were re-stimulated with S peptide pool. (**j-k**) Frequency of S protein-specific CD4^+^ T cells in circulation producing two (**j**) or three (**k**) of Th1 cytokines (IL-2, IFNγ, and TNFα) determined by intracellular cytokine staining (n = 3 per group). PBMCs were re-stimulated with S recombinant protein. * P<0.05, ** P<0.01, *** P<0.001, **** P<0.0001, ns: not significant.

PBMCs from baboons were collected from the blood after full schedule of vaccination (around day 80) and stimulated with S peptides pool (**Figure 4d; Figure S5a-b**) or S recombinant proteins (**Figure 4e-f; Figure S5c**). PBMCs from baboons immunized with 2.5 mg EV^SpikeM+P^ exhibited highest levels of IL-2, IFNγ, and TNFα secreting cells, with some variation across individual baboons (**Figure 4d-f; Figure S5**). Next, PBMCs were stimulated for ICCS analysis (**Figure S6**, S peptide stimulation (**Figure 4g-I; Figure S7**) and S recombinant protein stimulation (**Figure 4j-k; Figure S8**)). Increased expression of Th1 cytokines by CD4^+^ T was detected in the animals immunized with EV^SpikeM+P^ (**Figure 4g-k; Figure S7a; Figure S8a**). A trend towards increased IFNγ and TNFα secretion by CD8^+^ T cells was also observed, albeit was not statistically significant (**Figure S7b; Figure S8b**). Collectively, these results indicate that EV^SpikeM+P^ initiates humoral and cellular immune responses against the S protein antigen in a nonhuman primate model.

## Discussion

We report here a proof-of-concept study that generated engineered EVs that can delivery OVA mRNA and protein, and the S domain mRNA and protein of the SARS-CoV-2 virus to elicit functionally effective immunity. The engineered EVs vaccines, capsulated mRNA and protein induced immunity without overt toxicity. EVs are natural lipid nanovesicles and therefore are potentially better tolerated as an encapsulating carrier when compared to synthetic lipid nano-particles (LNPs). We show that administration of approximately 85% less OVA antigen with EV^OvaM+P^ compared to OVA^P^ (3 μg vs. 20 μg) resulted in more effective induction of T cell activation and control of tumor growth. While induction of humoral responses is critical for initial responses to foreign antigens in the context of infectious diseases, cellular immunity is thought to be critical for long-term protection (Wherry and Barouch 2022). In addition, many cancers are characterized by low T cell infiltration, which can prevent immune recognition and clearance of cancer cells and reduce the efficacy of immune checkpoint inhibitors (Ott and Wu 2019). This suggests that EV^X-M+P^ may result in enhanced antigen presentation to elicit T cell infiltration and activation to induce more durable long term memory responses compared to traditional protein vaccination. Additionally, CD47 on the surface of EVs may help to prevent phagocytosis by macrophages and increase accumulation in dendritic cells for peptide presentation, when compared to LNPs (Kamerkar, LeBleu et al. 2017). Such unique advantages of EVs compared to LNPs argue for its use in the healthy general population setting to prevent various diseases.

In addition, our simple EV based vaccine platform highlights many advantages that make it easy to deploy at a general population level for large scale studies to evaluate their efficacy and impact to prevent infectious diseases and malignant cancer. We specifically focused on methodologies that are easily reproducible with efficient and accessible technology in most laboratories with low cost and easy to execute EV vaccine manufacturing in the GMP facility. We predict that each dose of vaccine will cost less than $10 and can be easily shipped all over the world without the need for ultra-cold packaging. The use of both mRNA and protein make this vaccine platform more efficient. Collectively our study highlights the potential of EV^X-M+P^, as a viable platform for delivery of combination mRNA and protein antigens against infectious diseases and cancer.

## Acknowledgements

This work was supported by Lyda Hill Philanthropies^®^, Fifth Generation (Love, Tito’s), and Bosarge Family Trust to the Kalluri Laboratory at the University of Texas MD Anderson Cancer Center. The baboon studies were supported by NIH grant P40OD024628. The High Resolution Electron Microscopy Facility was supported by CCSG grant NIH P30CA016672. The Sanger Sequencing for vectors was supported by NCI grant CA016672 (SMF). We thank Jennifer Leveille, Laura Snowden, and Sarah J. Neal Webb for assistance with the study.

## Author Contributions

RK conceptually designed the strategy for this study, provided intellectual input and contributed to the writing of the manuscript. KMM contributed to the overall experimental design, the generation pseudovirus, and the writing and proofreading of the original manuscript draft. VSL contributed to the deisgn of EV^SpikeM+P^ and advised the design of the project in the early stage. XL generated cell lines and EVs, performed EVs characterization, prepared vaccines and performed vaccination, antibody detection, neutralization and lymphocytic cells isolation on mice for EV^SpikeM+P^. KAA and SJM generated the 293F-OVA EVs and performed EV characterization. KAA, SJM, and VB performed the OVA vaccination and evaluation on mice for EV^OvaM+P^. KAA conducted the OVA antibody detection. KAC performed bleeding of mice for ELISA. XL conducted the antibody detection, PBMCs isolation, ICCS, flow cytometry on baboons and all the data analysis, prepared the figures and wrote the original manuscript draft. YH performed splenocyte isolation from mice. SVK performed flow cytometry analysis. JH and DPD generated cell lines. JD contributed to the generation of pseudovirus. NTR and SS completed the mouse lymphocytic cells ICCS and flow cytometry experiment. JHS performed the vaccination of baboons. SC performed the ELISpot of baboon PBMCs and wrote the methods of ELISpot. MLK and HS performed mouse vaccination and bleeding for ELISA. KKM advised on the flow data analysis for EV^OvaM+P^.

## Figure legends

**Figure S1. Uncropped Western blots for Figure 1f-1g**. Uncropped Western blots of 293F OVA^+^ cells for OVA and vinculin (Figure 1f) (**a**), 293F OVA^+^ EVs for OVA and CD81 (Figure 1g) and quantification (n = 3) (**b**), and B16F10 OVA^+^ cells for OVA and vinculin (**Figure 1h**) (**c**). n.d., not detected.

**Figure S2. T cells proliferation in B16 tumors of vaccinated mice.** (**a**) Representative images of T cell proliferation by TSA staining in WT tumor and OVA tumor tissue from EV and Ova^P^ and EV^OvaM+P^ vaccinated mice. Scale bar, 100 μm. (**b**) Quantification of CD8^+^ T and proliferative CD8^+^ T cells (Ki67^+^) from TSA staining images.

**Figure S3. Uncropped Western blots for Figure 2e.** Uncropped Western blots for S protein (**a**), Syntenin-1 (**b**), and CD81 (**c**).

**Figure S4. Flow cytometry gating for the murine T cell phenotyping.**

**Figure S5. Baboon T cells phenotyping by ELISpot.** Quantification of cytokines (IL-2, IFNγ, and TNFα) releasing spots from S protein-specific T cells in circulation re-stimulated with S peptide pool (**a**) or recombinant S protein (**b-c**) by ELISpot (n = 3 per group). Data are shown as SFCs per 2 x 10^6^ cells.

**Figure S6. Flow cytometry gating for the baboon T cell phenotyping.**

**Figure S7. Baboon T cells activation by a S protein peptide pool.** Frequency of S protein-specific CD4^+^ (**a**) T cells and CD8^+^ T cells (**b**) in circulation producing single and multiple combinations of Th1 cytokines (IL-2, IFNγ, and TNFα) determined by intracellular cytokine staining (n = 3).

**Figure S8. Baboon T cells activation by a S protein peptide pool.** Frequency of S protein-specific CD4^+^ (**a**) T cells and CD8^+^ T cells (**b**) in circulation producing single and multiple combinations of Th1 cytokines (IL-2, IFNγ, and TNFα) determined by intracellular cytokine staining (n = 3).

## Materials and Methods

### Cell culture and transfection

FreeStyle^TM^ 293F (293F) cells from Gibco (R79007) were cultured in FreeStyle 293 expression media (Gibco) on a shaker with a speed of 35 rpm in a 37°C, 8% CO2 incubator. 293F cells were transfected with plasmids containing Spike protein (ATUM plasmid pD2528-CMV with insert QHD43416.1) and Spike-2P protein (site-directed mutagenesis to generate the 2P mutation on the plasmid containing Spike protein, GenScript) by 293fectin^TM^ transfection reagent (Gibco) under the selection of blasticidin (6 µg/mL, Gibco). 293T/17 (293T) cells from ATCC (CRL-11268), B16F10 cells from ATCC (CRL-6475), and Vero E6 TMPRSS2^+^ cells from Japanese Collection of Research Bioresources (JCRB) were cultured in complete DMEM (Dulbecco’s Modified Eagle’s Media (DMEM, Corning) containing 10% fetal bovine serum (FBS, Gemini), 1% penicillin-streptomycin (PS, Corning)) in a 37°C, 5% CO2 incubator. Vero E6 TMPRSS2^+^ cells were cultured under the selection of G-418 (1,000 µg/mL, Gibco). Cell viability and number were measured with trypan blue reagent with a Cellometer Mini (Nexcelom).

### Lentivirus generation and transduction

Lentivirus plasmids were created using Gateway cloning following manufacturer’s directions. The OVA entry fragment was created using a PCR amplicon from OVA plasmid (Addgene plasmid 64599) with primers containing attB sites (**Table 1**) and Phusion Plus DNA Polymerase (Thermo Fisher, F530N). OVA entry fragment was cloned into donor vector pDONR221 (Invitrogen, 12536017) utilizing BP Clonase II (Invitrogen, 11789020). Plasmids with correct insert were cloned into a lentivirus backbone destination vector (Addgene, plasmid 17451) with LR Clonase II Enzyme Mix (Invitrogen, 11791020). The lentivirus backbone plasmids with correct insert were verified with Sanger Sequencing.

**Table 1:**
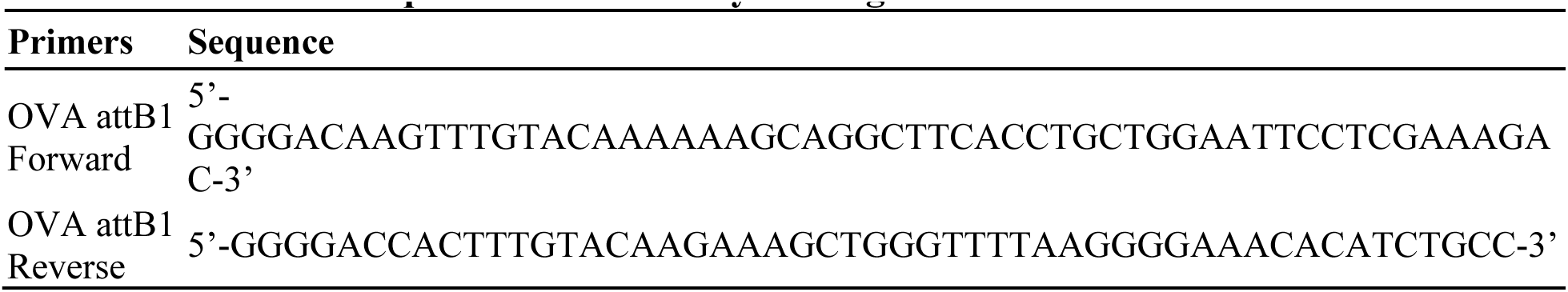
Ovalbumin sequences for Gateway cloning.

293F OVA^+^ cells and B16F10 OVA^+^ cells were created with lentiviral transduction. 293T cells were cultured at 37°C and 5% CO2 until they reached 70% confluency, followed by transfection with OVA lentivirus transfer vector and lentivirus packaging vectors psPAX2 (Addgene, plasmid 12260) and pMD2.G (Addgene, plasmid 12259) using Lipofectamine3000 transfection reagent (Invitrogen, L3000001) according to manufacturer’s directions. After 18 hours, 293T cell complete DMEM media was replaced with 10 mL of harvest media (30% FBS for complete media). Harvest media was then collected and concentrated using an Amicon Ultra-4 100 kD centrifugation filter (Amicon, UFC810024) at 1,500 x g for 30 minutes at room temperature. The concentrated harvest media was then added to 293F cells and B16F10 cells. After the treated 293F cells and B16F10 cells reached 1.5 x 10^6^ cells/mL and 90% confluence, respectively, blasticidin was added at a concentration of 6 μg/mL. After two weeks under selection, the cells were considered stable and confirmed by western blot. Blasticidin was maintained in the culture media throughout the course of cell culture.

### EVs isolation

293F cells were cultured in spinner flasks with initial concentration of 3 x 10^5^ cells/mL for 48-72 hours. Cells containing media were collected and centrifuged at 100 x g for 5 min to remove the cells and the supernatant was then centrifuged for 5 min at 800 x g followed by a second centrifugation for 10 min at 2,000 x g at room temperature. The supernatant was filtered through a 0.2 µm filter (Thermo Fisher) and then ultracentrifuged for 3 hours with a SW32 Ti rotor (Beckman Coulter) at 100,000 x g at 4°C. The pellet containing EVs was resuspended in 100 µl PBS (Corning) on a shaker at 4°C overnight followed by resuspension with a pipette for 1 minute. The size and concentration of EVs were measured using nanoparticle tracking analysis (NTA) with Malvern NanoSight NS300, according to the manufacturer’s directions. Isolated EVs were stored at -80°C until use. Cells were counted with a Cellometer Mini after the removal of conditioned media for the production of EVs and EV proteins from the cells.

### MicroBCA protein quantification

The protein concentration of EVs was measured using a Micro BCA^TM^ Protein Assay kit (Thermo Fisher) according to the manufacturer’s directions. Absorbance was read with FLUOstar Omega (BSI) at 562 nm.

### Transmission Electron Microscopy (TEM)

EVs isolated from cells were washed with ultracentrifugation for 3 hours with a SW41 Ti rotor (Beckman Coulter) at 100,000 x g at 4°C. The pellet was resuspended in PBS on a shaker overnight at 4°C followed by pipetting up and down for 1 minute. Around 3 x 10^9^ washed EVs were diluted in 10% glutaraldehyde (Electron Microscopy Sciences) to reach a final concentration of 2% glutaraldehyde. Samples were then placed to adhere on 100 mesh carbon coated, formvar coated copper grids treated with poly-l-lysine for 1 hour then negatively stained for 1 min with Millipore-filtered aqueous 2% uranyl acetate. The stain was blotted dry with the sample from the grids with filter paper. Samples were examined in a JEM 1010 transmission electron microscope (JEOL, USA, Inc., Peabody, MA) at an accelerating voltage of 80 kV. Digital images were obtained using the AMT Imaging System (Advanced Microscopy Techniques Corp., Danvers, MA).

### Flow cytometry analysis of EVs

5 x 10^9^ EVs, quantified by NTA, were resuspended in 200 μl of PBS and 10 µl of 4 µm aldehyde/sulfate latex (Invitrogen) beads were added per sample followed by a 15 min incubation at room temperature with rotation. 600 µl of PBS was then added to the EVs-beads mixture and incubated overnight at 4°C on a rotator. 400 µl of 1M glycine in PBS was added to each sample followed by a 30 min incubation with rotation at room temperature. The samples were then blocked with 100 µl of 10% BSA in PBS for 1 hour at room temperature. BSA was removed after centrifugation and the pellet was resuspended and incubated with 0.2 µg of primary antibodies (Mouse anti Human -isotype, -CD9, -CD47, -CD63, -CD81, **Table 2**) prepared in 20 µl of 2% BSA in PBS for 1 hour at room temperature with rotation. After three washes with 2% BSA in PBS, the samples were incubated with 1 µg of secondary antibodies (Donkey anti Mouse IgG Alexa Fluor 488, **Table 2**) in 20 µl of 2% BSA in PBS protected from light for 1 hour at room temperature with rotation. After three washes, the samples were resuspended in 300 µl of 2% BSA in PBS and analyzed on BD LSRFortessa^TM^ X-20. The data was analyzed using FlowJo (BD Biosciences).

**Table 2:** EV biomarker flow cytometry antibodies.

| Antibody | Vendor | Catalog number |
| --- | --- | --- |
| Isotype | BD Biosciences | 555746 |
| CD9 | Sigma Aldrich | SAB4700092 |
| CD47 | eBioscience | 14-0479-82 |
| CD63 | BD Bioscience | 556019 |
| CD81 | BD Biosciences | 555675 |
| Alexa Fluor 488 | Invitrogen | A21202 |

### Western blot analysis

Cells proteins were lysed and isolated from cell pellet by RIPA buffer (Thermo, 89901) with cOmplete mini EDTA free protease inhibitor (Roche, 11836170001) on ice incubation for 30 min and centrifugation at 15,000 rpm for 15 min at 4°C. Cell proteins and EVs proteins were quantified by Qubit Protein Assay (Invitrogen) using a Qubit 3.0 Fluorometer (Thermo Fisher) according to the manufacturer’s direction. 5-100 ng of S-RBD protein (SinoBiological, 40592-V08B) or ovalbumin (Sigma-Aldrich, A5503) recombinant protein were added as control. 20 μg cell proteins or 3-5 μg EVs proteins were denatured in LDS sample buffer containing 1M Dithiothreitol (DTT, Sigma) for 10 min at 70°C and loaded onto a 4-12% precast polyacrylamide gels for electrophoresis. The samples on the gel were then transferred to a methanol-activated PVDF membrane using a semi-dry transfer system (BioRad Trans-Blot Turbo Transfer). The membrane was blocked with 5% non-fat dry milk or 5% BSA in 1 x Tris-Buffered Saline with 0.1% Tween-20 (TBS-T) at room temperature on a shaker for 1 hour followed by an incubation of primary antibodies (**Table 3**) diluted in 2 or 5% BSA in TBS-T overnight on a shaker at 4°C or 5% BSA in TBS-T for ovalbumin blots. The membrane was then washed three times with TBS-T followed by an incubation with secondary antibodies (**Table 3**) diluted in 2% BSA in TBS-T or 5% BSA in TBS-T for ovalbumin blots for 1 hour on a shaker at room temperature. After three washes with TBS-T, the membrane was then incubated with West-Q Pico ECL reagent (genDEPOT) and chemiluminescence captured on a film (Cytiva).

**Table 3:**
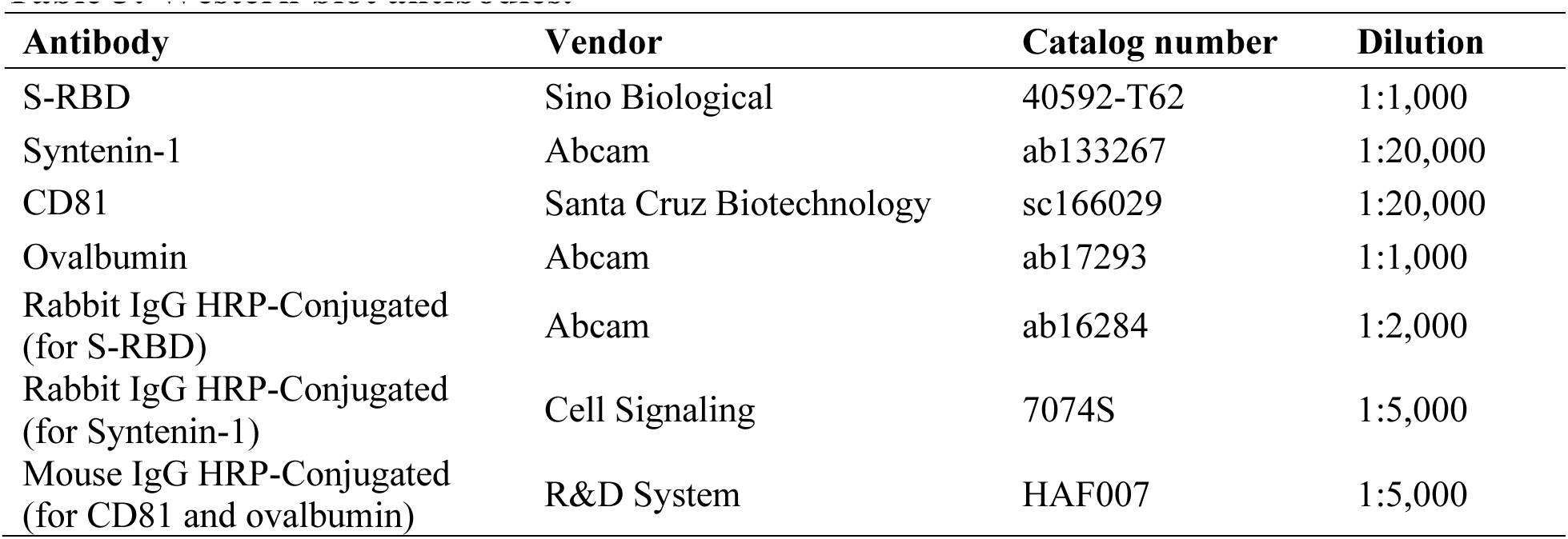
Western blot antibodies.

### qRT-PCR for the quantification of Spike and OVA mRNA from cells and EVs

mRNA was isolated from cells using RNeasy^®^ Mini Kit (Qiagen, Purification of Total RNA form Animal Cells Using Spin Technology) and EVs using Total Exosome RNA & Protein Isolation Kit (Invitrogen) following the manufacturer’s instructions and quantified by NanoDrop (Thermo). mRNA was then reverse transcribed into cDNA with High Capacity cDNA Reverse Transcription Kit (Invitrogen) with RNase Inhibitor (Invitrogen). qRT-PCR was performed with 20 ng of cDNA with PowerUp^TM^ SYBR^TM^ Green Master Mix (Applied Biosystems, A25742) and forward and reverse primers (**Table 4**).

**Table 4:**
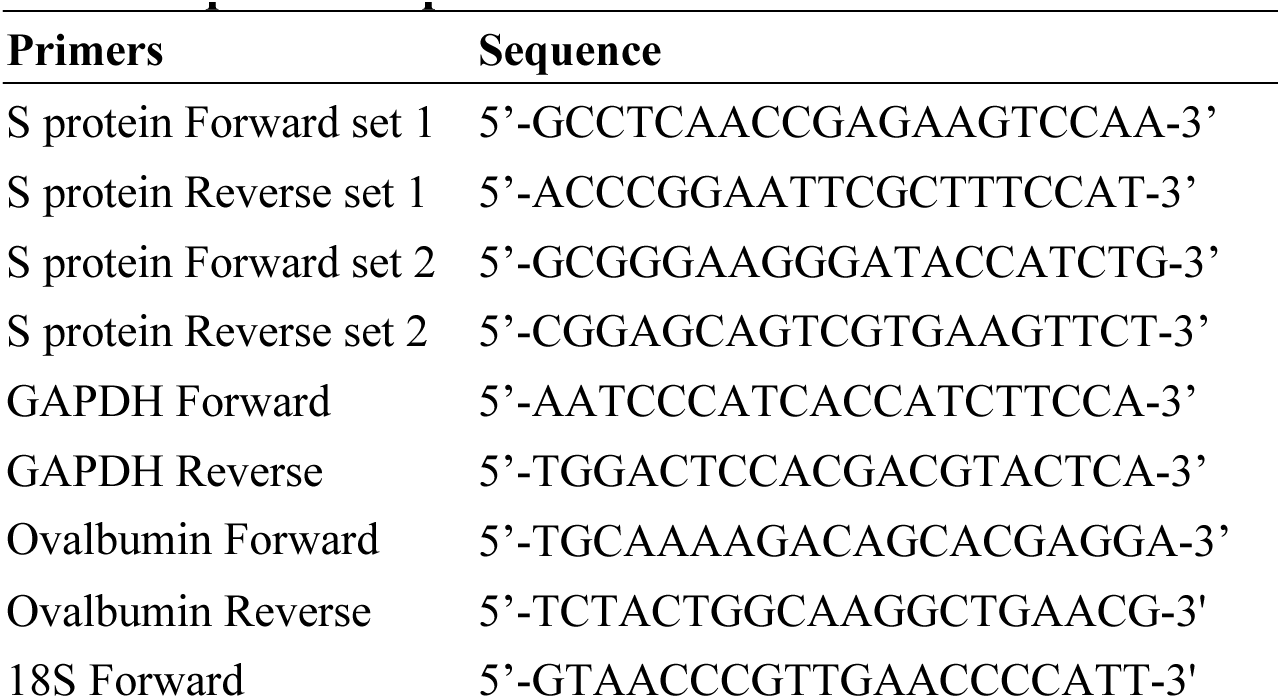
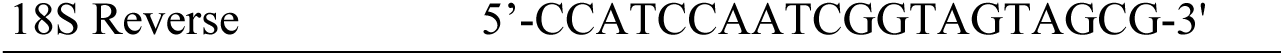
qRT-PCR primers.

### Murine vaccination and plasma collection

6 week old male and female Balb/c mice (Charles River Laboratories) or 6 week old male and female C57BL/6J mice (Jackson Laboratory) were injected with PBS, 100 μg wild type EVs, 100 μg EV-SC2S, 100 μg EV^SpikeM+P^ (1:1 mixed with Alhydrogel, InvivoGen, vac-alu-250) intramuscularly in the quadriceps (day 0), with two boosters on day 14 and day 28. The mice were euthanized on day 42. The body weight of the mice was measured weekly. Approximately 100 μl blood was collected using the retro-orbital plexus at different time points and mixed with anticoagulant (heparin) immediately. Plasma was collected after centrifugation at 6,000 rpm at 4°C for 5 min and stored at -80°C for future use. Gender balance was maintained with experimental mice and all experiments were approved by the Institutional Animal Care and Use Committee at MD Anderson Cancer Center.

For tumor challenge experiments, mice were vaccinated with PBS, 100 μg wild type EVs, 100 μg EV-SC2S, 100 μg EV^OvaM+P^, 20 μg of Ovalbumin recombinant protein (Sigma-Aldrich, A5503) (1:1 mixed with Alhydrogel) intramuscularly in the quadriceps at day 0, with boosters at day 14 and day 28. Blood was collected from the facial vein every two weeks for the first 28 days and at endpoint. At day 42, mice were challenged with subcutaneous tumors. To test antigen specificity, 1x10^6^ B16F10 wide type cells were injected on the right flank and 1x10^6^ B16F10 OVA cells were injected on the left flank. Mice were monitored and weighed every other day to track tumor growth. Mice were then euthanized prior to reaching moribundity, as determined by tumor ulceration or tumor diameter exceeding 2.0 cm.

### Baboon vaccination and plasma collection

8-25 kg male and female Olive baboons were injected with PBS, 0.625 mg EV^SpikeM+P^, 1.25 mg EV^SpikeM+P^, 2.5 mg EV^SpikeM+P^ (1:1 mixed with Alhydrogel) intramuscularly (day 0), with two boosters on day 21 and day 42. 30 mL blood was collected with anticoagulant (K2EDTA) at different time points post vaccination until around 6 months. Plasma was collected after centrifugation at 4,000 rpm at 4°C for 10 min and stored at -80°C for future use. PBMCs were isolated and stimulated with vaccine protein or peptides to monitor antigen specific T cell immune responses as described below. All experiments were approved by the Institutional Animal Care and Use Committee at MD Anderson Cancer Center.

### ELISA

For the detection of anti-S-RBD IgG antibodies from plasma, 50 ng S-RBD recombinant protein (Sino Biologicals) in coating buffer (15mM sodium carbonate, 35mM sodium bicarbonate in Milli-Q water, pH 9.6) was added to MaxiSorp C-shaped 96 well plates (Thermo Scientific), sealed to precent evaporation, at 4°C overnight. For the detection of anti-Ovalbumin IgG antibodies from plasma, 200 μg of Ovalbumin recombinant protein in coating buffer (15mM sodium carbonate, 35mM sodium bicarbonate in Milli-Q water, pH 9.6) was added to MaxiSorp C-shaped 96 well plates (Thermo Scientific), sealed to prevent evaporation, at 4°C overnight. A no antigen control was included, where wells were coated with coating buffer alone without assay specific recombinant protein. The coating substrate was removed from the plate the next day. The plate was then blocked with 5% non-fat milk in TBS for 1 hour at room temperature followed by 1 hour of incubation of the plasma diluted 1:50 in TBS, at 37°C. The plate was washed three times with TBS-T, then incubated with goat anti-mouse secondary HRP antibody (Sigma-Aldrich, #A0168, 1:10,000, for mouse plasma) or goat anti-human secondary HRP antibody (Sigma-Aldrich, #A0170, 1:10,000, for baboon plasma) in 1% BSA in TBS for 1 hour at room 37°C. After three washes with TBS-T, 50 μl of 1-Step^TM^ Ultra TMB-ELISA (Thermo) was added to the plate. 50 μl of TMB stopping reagent (0.18M sulfuric acid) was then added to stop the reaction after 15 min and the absorbance at 450 nm was read immediately using the VersaMax Microplate Reader. For the titration of anti-S-RBD IgG antibodies from vaccinated baboons, the plasma was serially diluted. Data were presented as the absorbance of wells with no antigen (diluted plasma) subtracted from the absorbance of wells containing antigen (McAndrews, Dowlatshahi et al. 2020).

### Neutralization Assay

8 x 10^5^ 293T cells were seeded in 6-well plate and transfected with pcDNA3-FLAG-VSVG plasmids (Addgene, plasmid 80606) for 24 hours with 50 μl of purchased or previously collected VSVΔG-Luc pseudovirus (Kerafast) using Lipofectamine 3000 (Invitrogen) according to the manufacturer’s directions. The cells were washed three times with 1% FBS in PBS after 24 hours and fresh media was added. Conditioned media containing virions was collected in 24 hours and centrifuged at 1,300 rpm for 5 min, followed by 0.45 μm filtration (Corning). The viruses were titered with Vero E6 TMPRSS2^+^ cells and stored at -80°C for future use. To titer the virus, 1 x 10^5^ Vero E6 TMPRSS2^+^ cells/mL were seeded in 96-well plate. Serially diluted virus was added to the cells 24 hours later. The luciferase signals were measured using the Luciferase Reporter Gene Assay System (Invitrogen) and FLUOstar Omega (BSI). VSVΔG-Luc-Spike pesudovirus was generated and titered in a similar manner with the pcDNA3-FLAG-VSVG plasmid replaced with plasmids containing S protein. For the neutralization assay, the virus was incubated with plasma in 1:20 dilution at 37°C for 1 hour before adding to the cells. Data are reported as raw relative luminescence units (RLU).

### ELISpot assay

Peripheral blood mononuclear cells (PBMCs) were isolated fresh from blood using Ficoll-Paque Plus (Cytvia) gradient. PBMC collected were washed twice with 1x PBS followed by complete RPMI (Roswell Park Memorial Institute 1640 medium (RPMI, Corning) containing 10% FBS, glutamine and 1% PS). Cells were counted (Cellometer Auto 2000; Nexcelom) and adjusted to numbers needed for ELISpot assay. IL-2, IFNγ and TNFα ELISpot were performed using monkey cytokine specific ELISpotPLUS (ALP) KITs (Mabtech, Inc.) following the manufacturer’s instructions. Assays were performed in complete RPMI medium. PepTivator^®^ SARS-CoV-2 Prot_S-peptide pool (Miltenyi Biotec, consisting of 15-mer sequences with 11 amino acids overlap covering the complete protein coding sequence (aa 5–1273) of S protein) and recombinant S protein (Sino Biological) were used for stimulation at a final concentration of 2 μg/mL; Con A (Sigma) at same concentration and unstimulated PBMCs in medium alone, were used as positive and negative controls respectively. Freshly prepared PBMCs (2 × 10^5^ per well) in duplicates were added with peptide, protein or controls and incubated at 37 °C with 5 % CO2 for ∼36-40 hours, without disturbing the plate during incubation. Activated cytokine secreting cells as positive spots were detected following the manufacturer’s protocol. Plates were completely dried and scanned using the Mabtech IRISTM FluoroSpot/ELISpot Reader (Sweden), for counting, imaging and analysis.

### Surface and intracellular cytokine staining

For murine cells surface and intracellular cytokine staining (ICCS), murine spleens and lymph nodes were dissociated through 70 μm mesh strainers followed by red blood cell lysis (Life Technologies, A1049201). Tumors were minced and digested with 1.5 mg/mL collagenase type I (Sigma-Aldrich, SCR103) in PBS at 37°C and 150 rpm for 18 minutes, then dissociated through 70 μm mesh strainers with 5 mL complete RPMI (RPMI 1640 Medium with L-Glutamine, Corning,), 10mM sodium pyruvate (Corning), 1% PS and 10% heat-inactivated FBS). PBMCs were isolated by Ficoll-Paque Plus (Cytiva) from fresh whole blood with anticoagulant (heparin, Sigma-Aldrich). Blood was diluted in PBS and added on the top of Ficoll-Paque Plus (Cytiva) for a 40 min centrifuge at room temperature for 400 x g with no brake. The lymphocyte layer was isolated and washed with PBS at 5 min 400 x g twice. To detect S protein specific immune response, cells were stimulated with either PepTivator^®^ SARS-CoV-2 Prot_S-peptide pool (Miltenyi Biotec) or S recombinant protein (Sino Biological) at a final concentration of 1 μg/mL for 6 hours incubation in a 37 °C, 5 % CO2 incubator with protein transport inhibitor (Invitrogen) added in the last 4 hours. Cells were washed in 2 μM EDTA/PBS followed by centrifugation at 1,700 x rpm for 3 minutes and the supernatant removed. Cells were incubated with fixable viability dye (eBioscience) for 10-15 minutes followed by two washes with 2 μM EDTA/PBS. 4% paraformaldehyde (PFA) was applied to fix the cells for 10 minutes followed by two washes. Cells were incubated in BD Perm/Wash^TM^ buffer for 15 minutes then with intracellular antibodies for 2 hours. After two washes, cells were resuspended in 2 μM EDTA/2 % FBS/PBS for analysis on a BD FACSymphony^TM^ A3. Antibodies and fixable viability dye used are listed in **Table 5**.

**Table 5:**
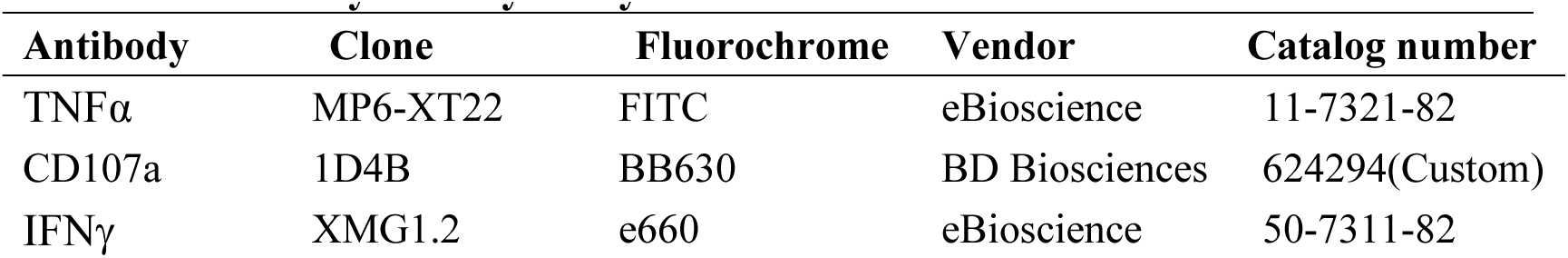
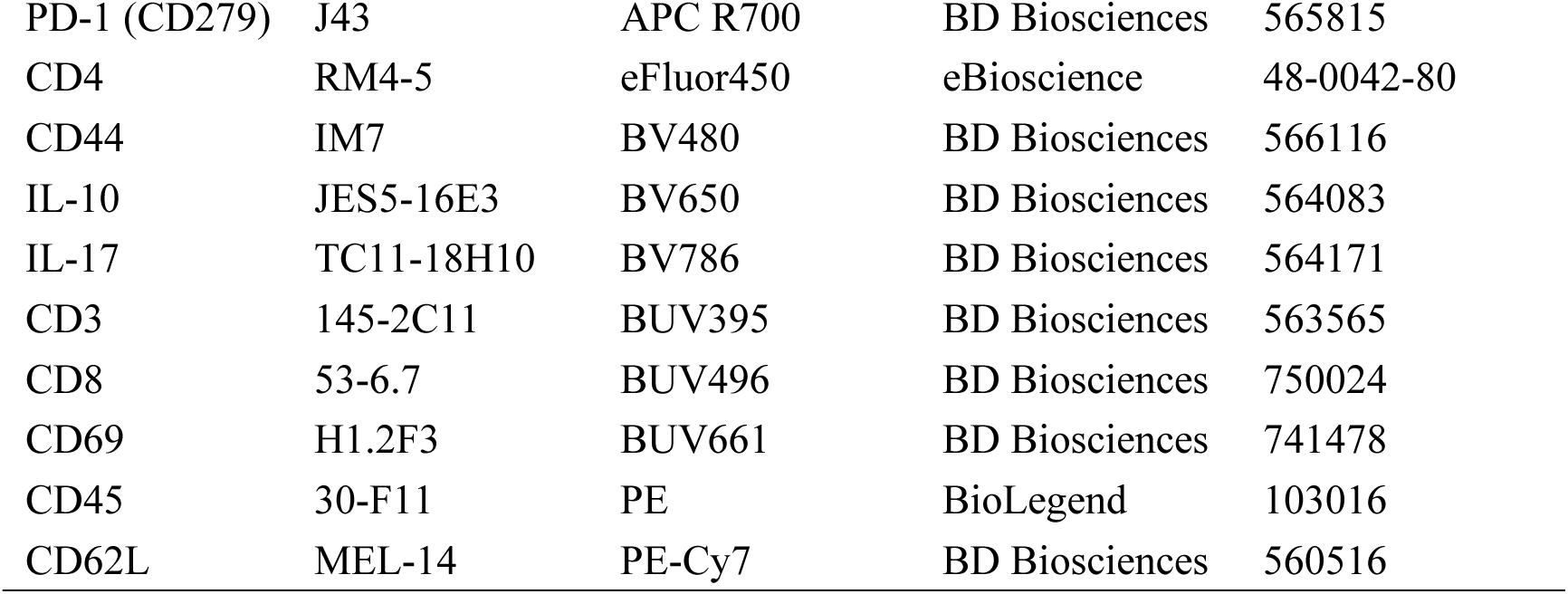
Anti-murine surface and intracellular cytokines antibodies for S protein/peptide stimulation flow cytometry analysis.

For staining of tumors, spleens, and lymph nodes of mice vaccinated for tumor challenge experiments, cells were incubated with 100 μL of antibody cocktail (surface antibodies, CD16/CD32 (BD), and fixable viability dye (eBiosciences, eFluor780)) on ice for 1 hour. After incubation, cells were centrifuged at 800 x g and 4°C for 3 minutes. Supernatant was aspirated, and cells were resuspended in 200 μL of FACS buffer (2% FBS/PBS). The cells were centrifuged and washed two more times. After final wash, supernatant was aspirated, and cells were resuspended in 200 μL of BD Cytofix and incubated on ice for 1 hour. After incubation, cells were spun down at 800 x g and 4°C for 3 minutes. Supernatant was aspirated, and cells were resuspended in 200 μL of FACS buffer. The cells were washed two more times, resuspended in 200 μL of FACS buffer, and stored at 4°C until analyzed. Flow cytometry analysis was performed with BD LSRFortessa^TM^ X-20 and FlowJo (BD Biosciences). Antibodies and fixable viability dye used are listed in **Table 6**.

**Table 6:**
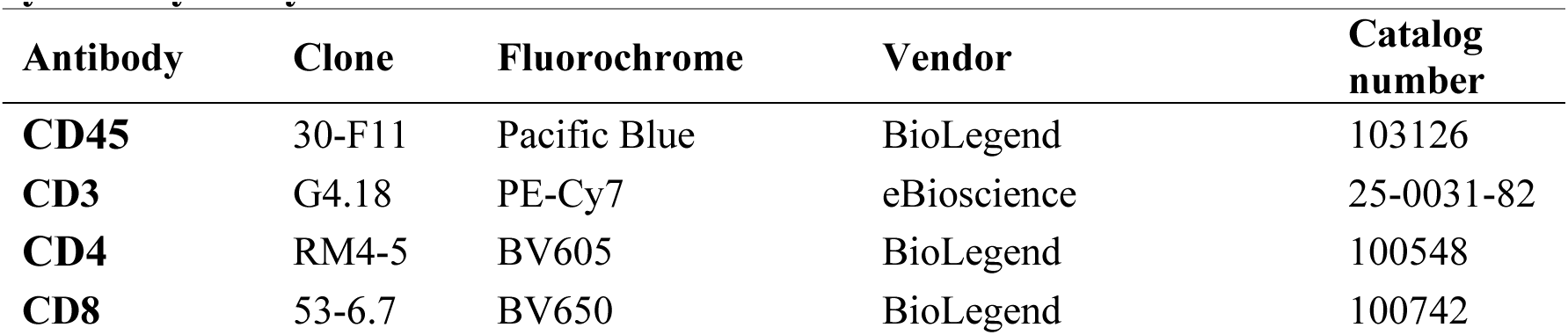
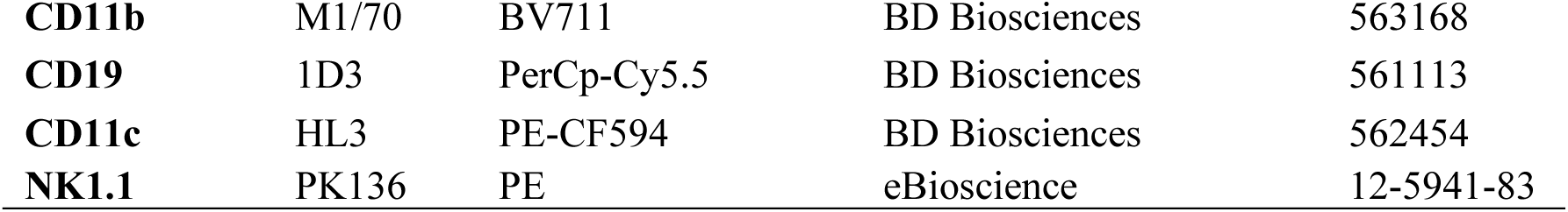
Anti-murine surface and intracellular cytokines antibodies for tumor challenge flow cytometry analysis.

For ICCS of baboon cells, Baboon PBMCs were isolated from fresh whole blood with anticoagulant (K2EDTA) and stimulated as described above. Cells were washed in FACS buffer with centrifugation at 400 x g for 5 minutes and remove the supernatant. Cells were then incubated with surface antibodies and LIVE/DEAD fixable yellow dye (Invitrogen) for 20 minutes followed two washes. BD Cytofix/Cytoperm^TM^ was used to fix and permeabilize the cells for 30 minutes followed by two washes. Cells were then incubated with intracellular antibodies for 30 minutes with two washes. Cells were resuspended in FACS buffer for analysis on BD LSRFortessa^TM^ X-20. Antibodies and yellow live/dead dye used are listed in **Table 7**. The data was analyzed using FlowJo (BD Biosciences).

**Table 7:** Anti-NHP surface and intracellular cytokines antibodies for flow cytometry analysis.

| **Antibody** | **Clone** | **Fluorochrome** | **Vendor** | **Catalog number** |
| --- | --- | --- | --- | --- |
| CD3 | SP34-2 | BV650 | BD Biosciences | 563916 |
| CD4 | L200 | APC-H7 | BD Biosciences | 560837 |
| CD8 | RPA-T8 | APC | BD Biosciences | 561421 |
| IL-2 | MQ1-17H12 | BV421 | BD Biosciences | 564164 |
| IFNγ | 4S.B3 | PerCP-Cy5.5 | BD Biosciences | 560742 |
| TNFα | A20 | PE-Cy7 | BD Biosciences | 560678 |

### TSA staining

3 to 5 <u>μ</u>m thick tissue sections were deparaffinized and rehydrated. Antigen retrieval was performed in TE buffer (pH 9.0) in a microwave for 15 minutes. Tissues were blocked for 10 minutes at room temperature with 1% BSA in TBS, then incubated with CD4 primary antibody for 1 hour at room temperature (**Table 8**). Afterwards, slides were washed 3 times with TBS for 2 minutes. Secondary (**Table 8**) was applied and incubated for 10 minutes at room temperature, then washed 3 times with TBS for 2 minutes. Slides were then incubated with TSA Plus (Opal 570, 1:100, Akoya Biosciences) for 10 minutes at room temperature, and immediately washed with TBS two times upon removal. Antigen retrieval was again performed in TE buffer (pH 9.0) in a microwave for 15 minutes. Slides were washed with dH2O for 2 minutes, followed by another 2-minute wash with TBS. Steps were repeated from the initial block to antigen retrieval to add Ki-67 primary antibody (**Table 8**), except with overnight incubation with the primary antibody at 4°C. Steps were again repeated from the initial block to antigen retrieval to add CD8a primary antibody (**Table 8**) with 1 hour incubation with the primary antibody at room temperature. Hoechst (Invitrogen) staining was applied at a concentration of 1:10,000 for 10 minutes, followed by two washes with TBS for 2 minutes. Slides were mounted with Vectashield (VectorLabs) and imaged using a BD-X Series Keyence microscope. Images were quantified using ImageJ.

**Table 8:** TSA staining antibodies.

| Antibody | Vendor | Catalog number | Dilution |
| --- | --- | --- | --- |
| CD4 | Cell Signaling | 25229S | 1:100 |
| Ki67 | Thermo Fisher | RM9106-S | 1:100 |
| CD8a | Cell Signaling | 98941S | 1:800 |
| Rabbit on Rodent HRP-Polymer | BioCare Medical | RMR622 | - |

## References

Beck, L. and H. L. Spiegelberg (1989). “The polyclonal and antigen-specific IgE and IgG subclass response of mice injected with ovalbumin in alum or complete Freund’s adjuvant.” Cell Immunol 123(1): 1–8.

Cheng, K. and R. Kalluri (2023). “Guidelines for clinical translation and commercialization of extracellular vesicles and exosomes based therapeutics.” Extracellular Vesicle 2: 100029.

Corbett, K. S., D. K. Edwards, S. R. Leist, O. M. Abiona, S. Boyoglu-Barnum, R. A. Gillespie, S. Himansu, A. Schäfer, C. T. Ziwawo and A. T. DiPiazza (2020). “SARS-CoV-2 mRNA vaccine design enabled by prototype pathogen preparedness.” Nature 586(7830): 567–571.

Dudani, R., Y. Chapdelaine, H. Faassen Hv, D. K. Smith, H. Shen, L. Krishnan and S. Sad (2002). “Multiple mechanisms compensate to enhance tumor-protective CD8(+) T cell response in the long-term despite poor CD8(+) T cell priming initially: comparison between an acute versus a chronic intracellular bacterium expressing a model antigen.” J Immunol 168(11): 5737–5745.

El-Shennawy, L., A. D. Hoffmann, N. K. Dashzeveg, K. M. McAndrews, P. J. Mehl, D. Cornish, Z. Yu, V. L. Tokars, V. Nicolaescu and A. Tomatsidou (2022). “Circulating ACE2-expressing extracellular vesicles block broad strains of SARS-CoV-2.” Nature communications 13(1): 405.

Escudier, B., T. Dorval, N. Chaput, F. André, M.-P. Caby, S. Novault, C. Flament, C. Leboulaire, C. Borg and S. Amigorena (2005). “Vaccination of metastatic melanoma patients with autologous dendritic cell (DC) derived-exosomes: results of thefirst phase I clinical trial.” Journal of translational medicine 3(1): 1–13.

Haltom, A. R., W. E. Hassen, J. Hensel, J. Kim, H. Sugimoto, B. Li, K. M. McAndrews, M. R. Conner, M. L. Kirtley and X. Luo (2022). “Engineered exosomes targeting MYC reverse the proneural-mesenchymal transition and extend survival of glioblastoma.” Extracellular Vesicle 1: 100014.

Harrison, A. G., T. Lin and P. Wang (2020). “Mechanisms of SARS-CoV-2 transmission and pathogenesis.” Trends in immunology 41(12): 1100–1115.

Heidelberger, M., K. O. Pedersen and A. Tiselius (1936). “Ultracentrifugal and Electrophoretic Studies on Antibodies.” Nature 138(3482): 165–165.

Hopkins, F. G. (1900). “On the separation of a pure albumin from egg-white.” J Physiol 25(4): 306–330.

Huntington, J. A. and P. E. Stein (2001). “Structure and properties of ovalbumin.” J Chromatogr B Biomed Sci Appl 756(1-2): 189–198.

Kalluri, R. and V. S. LeBleu (2020). “The biology, function, and biomedical applications of exosomes.” Science 367(6478).

Kalluri, R. and K. M. McAndrews (2023). “The role of extracellular vesicles in cancer.” Cell 186(8): 1610–1626.

Kamerkar, S., V. S. LeBleu, H. Sugimoto, S. Yang, C. F. Ruivo, S. A. Melo, J. J. Lee and R. Kalluri (2017). “Exosomes facilitate therapeutic targeting of oncogenic KRAS in pancreatic cancer.” Nature 546(7659): 498–503.

Ke, Y., Y. Li and J. A. Kapp (1995). “Ovalbumin injected with complete Freund’s adjuvant stimulates cytolytic responses.” Eur J Immunol 25(2): 549–553.

Lee, P., C.-U. Kim, S. H. Seo and D.-J. Kim (2021). “Current status of COVID-19 vaccine development: focusing on antigen design and clinical trials on later stages.” Immune network 21(1).

Lener, T., M. Gimona, L. Aigner, V. Börger, E. Buzas, G. Camussi, N. Chaput, D. Chatterjee, F. A. Court and H. A. d. Portillo (2015). “Applying extracellular vesicles based therapeutics in clinical trials–an ISEV position paper.” Journal of extracellular vesicles 4(1): 30087.

Li, Z., Z. Wang, P.-U. C. Dinh, D. Zhu, K. D. Popowski, H. Lutz, S. Hu, M. G. Lewis, A. Cook and H. Andersen (2021). “Cell-mimicking nanodecoys neutralize SARS-CoV-2 and mitigate lung injury in a non-human primate model of COVID-19.” Nature nanotechnology 16(8): 942–951.

Liu, H., Z. Xie and M. Zheng (2022). “Unprecedented chiral nanovaccines for significantly enhanced cancer immunotherapy.” ACS Applied Materials & Interfaces 14(35): 39858–39865.

McAndrews, K. M., D. P. Dowlatshahi, J. Dai, L. M. Becker, J. Hensel, L. M. Snowden, J. M. Leveille, M. R. Brunner, K. W. Holden and N. S. Hopkins (2020). “Heterogeneous antibodies against SARS-CoV-2 spike receptor binding domain and nucleocapsid with implications for COVID-19 immunity.” JCI insight 5(18).

Mendt, M., S. Kamerkar, H. Sugimoto, K. M. McAndrews, C.-C. Wu, M. Gagea, S. Yang, E. V. R. Blanko, Q. Peng and X. Ma (2018). “Generation and testing of clinical-grade exosomes for pancreatic cancer.” JCI insight 3(8).

Morse, M. A., J. Garst, T. Osada, S. Khan, A. Hobeika, T. M. Clay, N. Valente, R. Shreeniwas, M. A. Sutton and A. Delcayre (2005). “A phase I study of dexosome immunotherapy in patients with advanced non-small cell lung cancer.” Journal of translational medicine 3(1): 1–8.

Naqvi, A. A. T., K. Fatima, T. Mohammad, U. Fatima, I. K. Singh, A. Singh, S. M. Atif, G. Hariprasad, G. M. Hasan and M. I. Hassan (2020). “Insights into SARS-CoV-2 genome, structure, evolution, pathogenesis and therapies: Structural genomics approach.” Biochimica et Biophysica Acta (BBA)-Molecular Basis of Disease 1866(10): 165878.

Ohno, S.-i., M. Takanashi, K. Sudo, S. Ueda, A. Ishikawa, N. Matsuyama, K. Fujita, T. Mizutani, T. Ohgi and T. Ochiya (2013). “Systemically injected exosomes targeted to EGFR deliver antitumor microRNA to breast cancer cells.” Molecular Therapy 21(1): 185–191.

Ott, P. A. and C. J. Wu (2019). “Cancer Vaccines: Steering T Cells Down the Right Path to Eradicate Tumors.” Cancer Discov 9(4): 476–481.

Pallesen, J., N. Wang, K. S. Corbett, D. Wrapp, R. N. Kirchdoerfer, H. L. Turner, C. A. Cottrell, M. M. Becker, L. Wang and W. Shi (2017). “Immunogenicity and structures of a rationally designed prefusion MERS-CoV spike antigen.” Proceedings of the National Academy of Sciences 114(35): E7348–E7357.

Rotzschke, O., K. Falk, S. Stevanovic, G. Jung, P. Walden and H. G. Rammensee (1991). “Exact prediction of a natural T cell epitope.” Eur J Immunol 21(11): 2891–2894.

Van Niel, G., G. d’Angelo and G. Raposo (2018). “Shedding light on the cell biology of extracellular vesicles.” Nature reviews Molecular cell biology 19(4): 213–228.

Wang, Q., Y. Zhang, L. Wu, S. Niu, C. Song, Z. Zhang, G. Lu, C. Qiao, Y. Hu and K.-Y. Yuen (2020). “Structural and functional basis of SARS-CoV-2 entry by using human ACE2.” Cell 181(4): 894–904. e899.

Wherry, E. J. and D. H. Barouch (2022). “T cell immunity to COVID-19 vaccines.” Science 377(6608): 821–822.

Wu, J. T., K. Leung and G. M. Leung (2020). “Nowcasting and forecasting the potential domestic and international spread of the 2019-nCoV outbreak originating in Wuhan, China: a modelling study.” The Lancet 395(10225): 689–697.

Xu, J., J. Lv, Q. Zhuang, Z. Yang, Z. Cao, L. Xu, P. Pei, C. Wang, H. Wu and Z. Dong (2020). “A general strategy towards personalized nanovaccines based on fluoropolymers for post-surgical cancer immunotherapy.” Nature Nanotechnology 15(12): 1043–1052.

Yáñez-Mó, M., P. R.-M. Siljander, Z. Andreu, A. Bedina Zavec, F. E. Borràs, E. I. Buzas, K. Buzas, E. Casal, F. Cappello and J. Carvalho (2015). “Biological properties of extracellular vesicles and their physiological functions.” Journal of extracellular vesicles 4(1): 27066.

