## Supplementary figures and images for "Development of an engineered extracellular vesicles-based vaccine platform for combined delivery of mRNA and protein to induce functional immunity"

### Supplemental figures

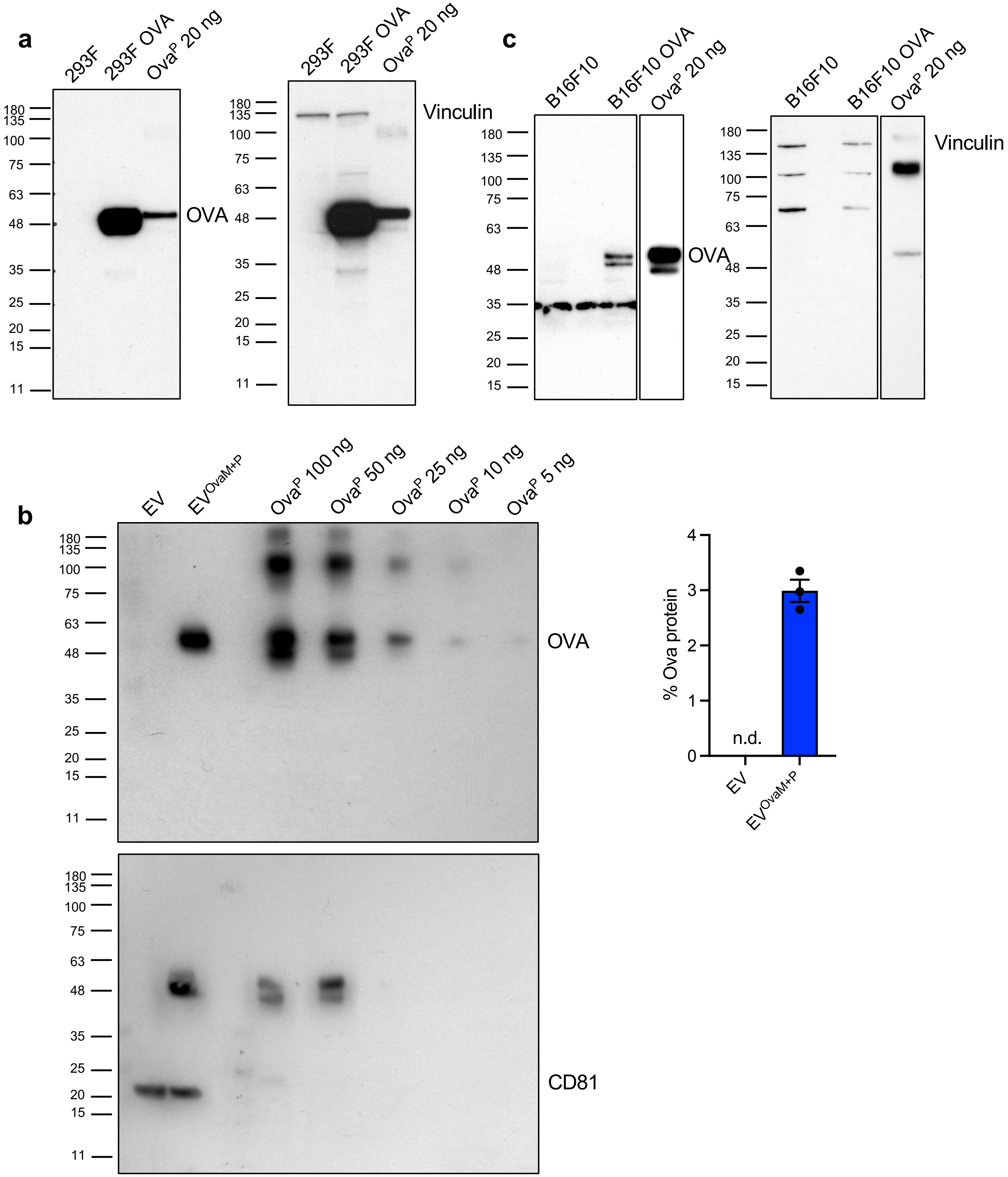

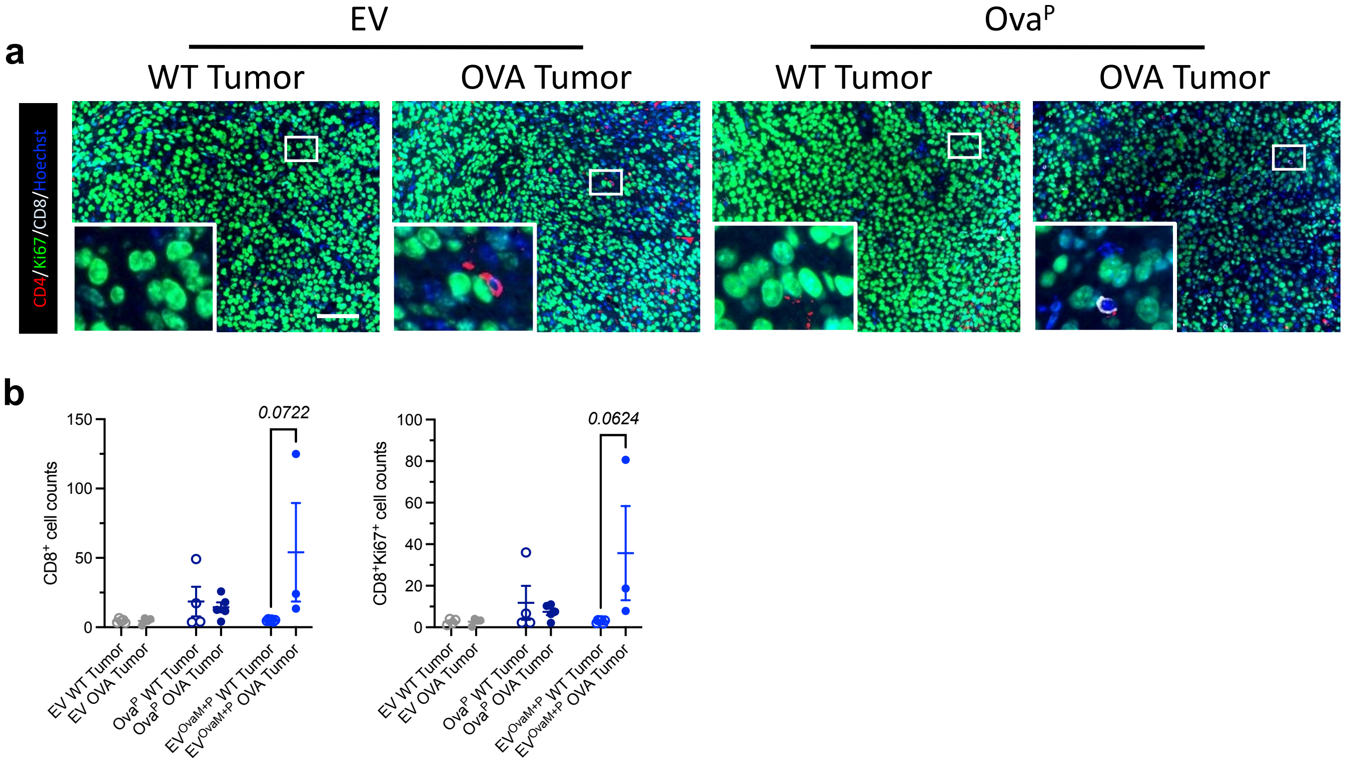

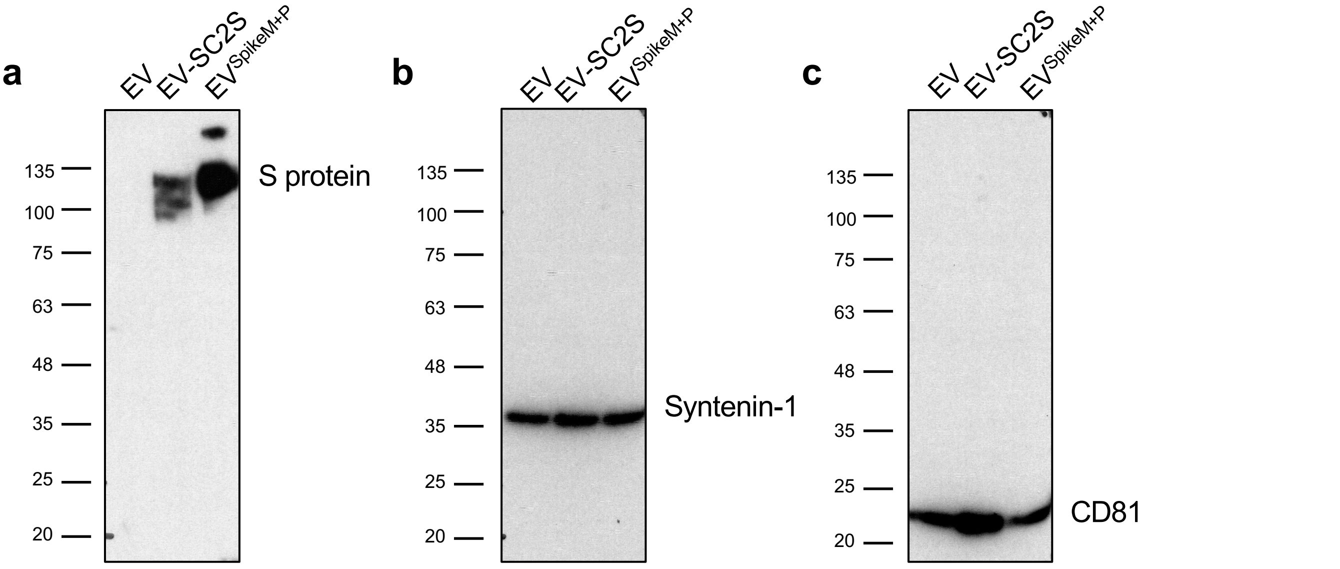

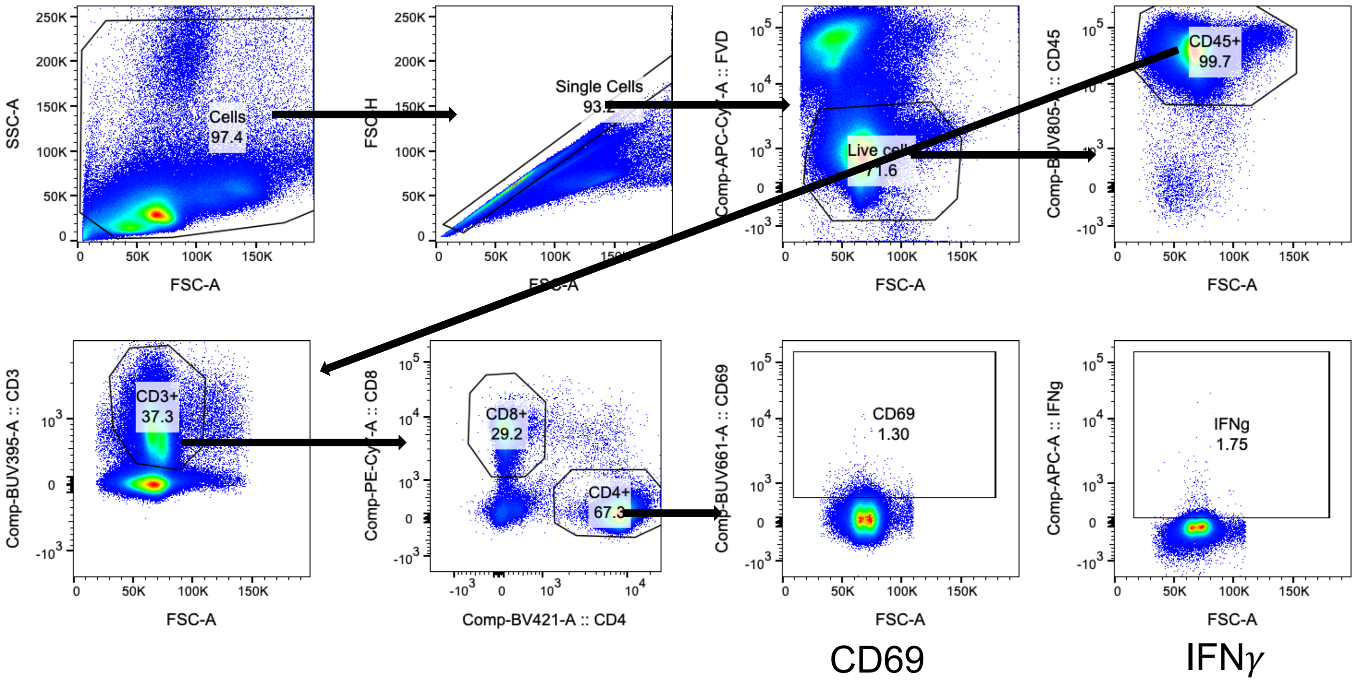

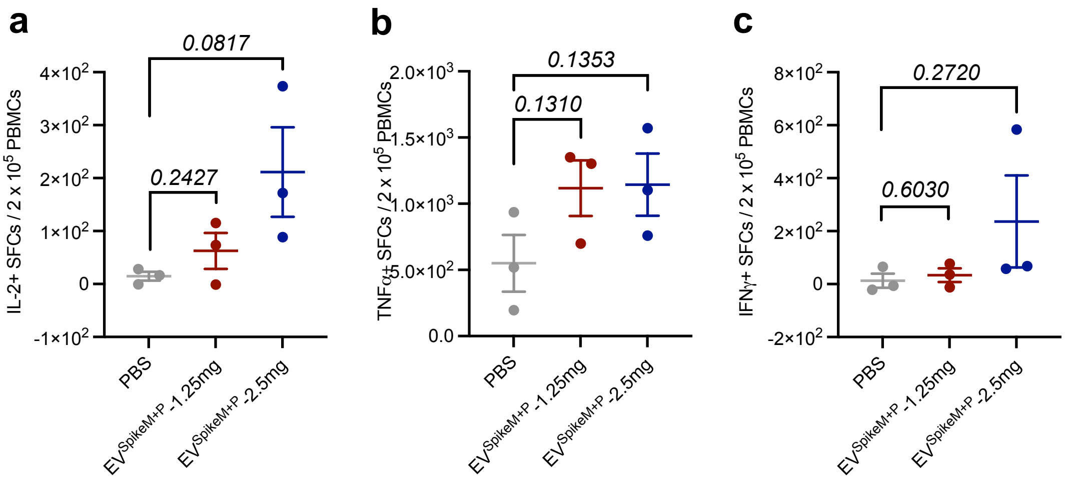

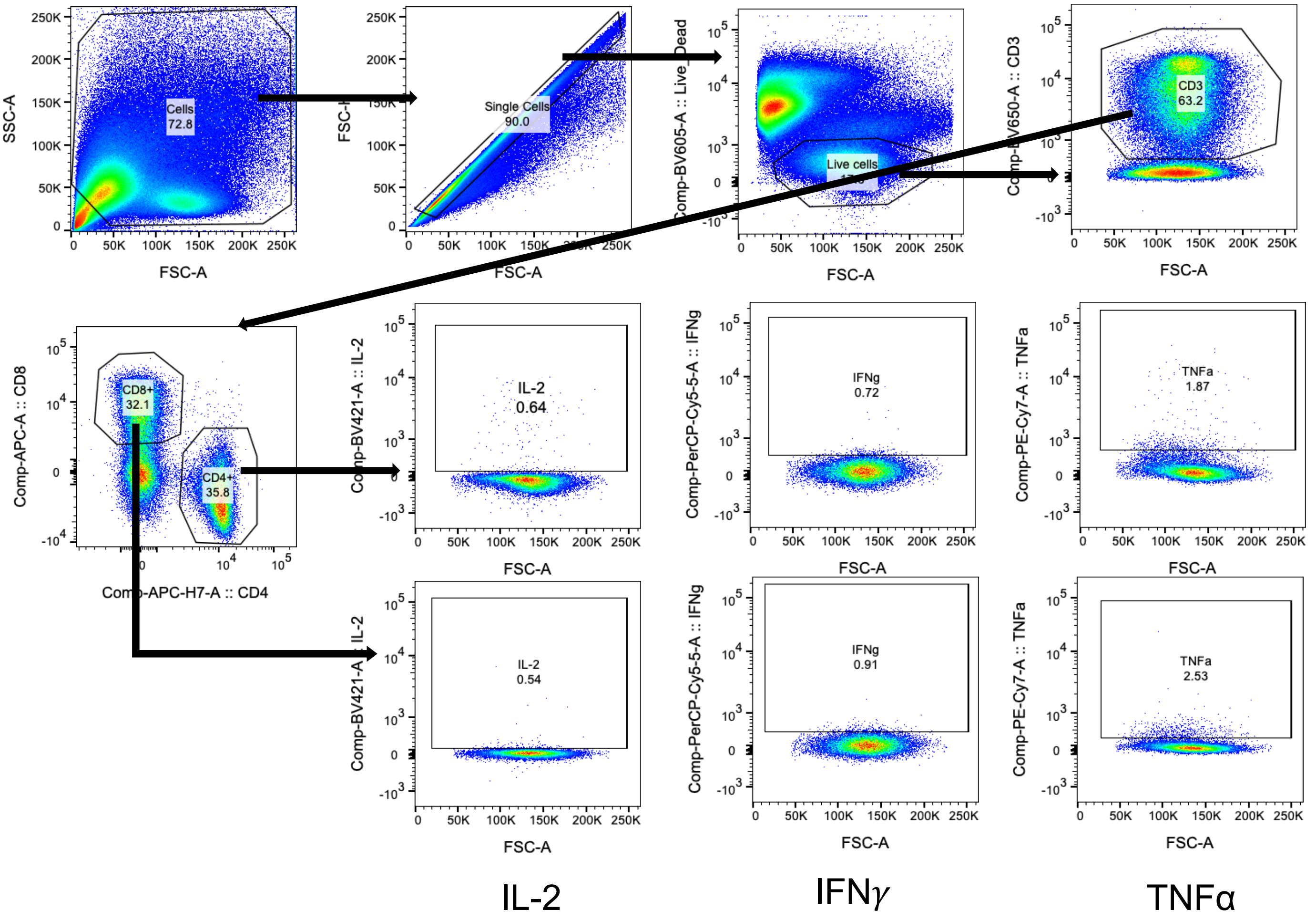

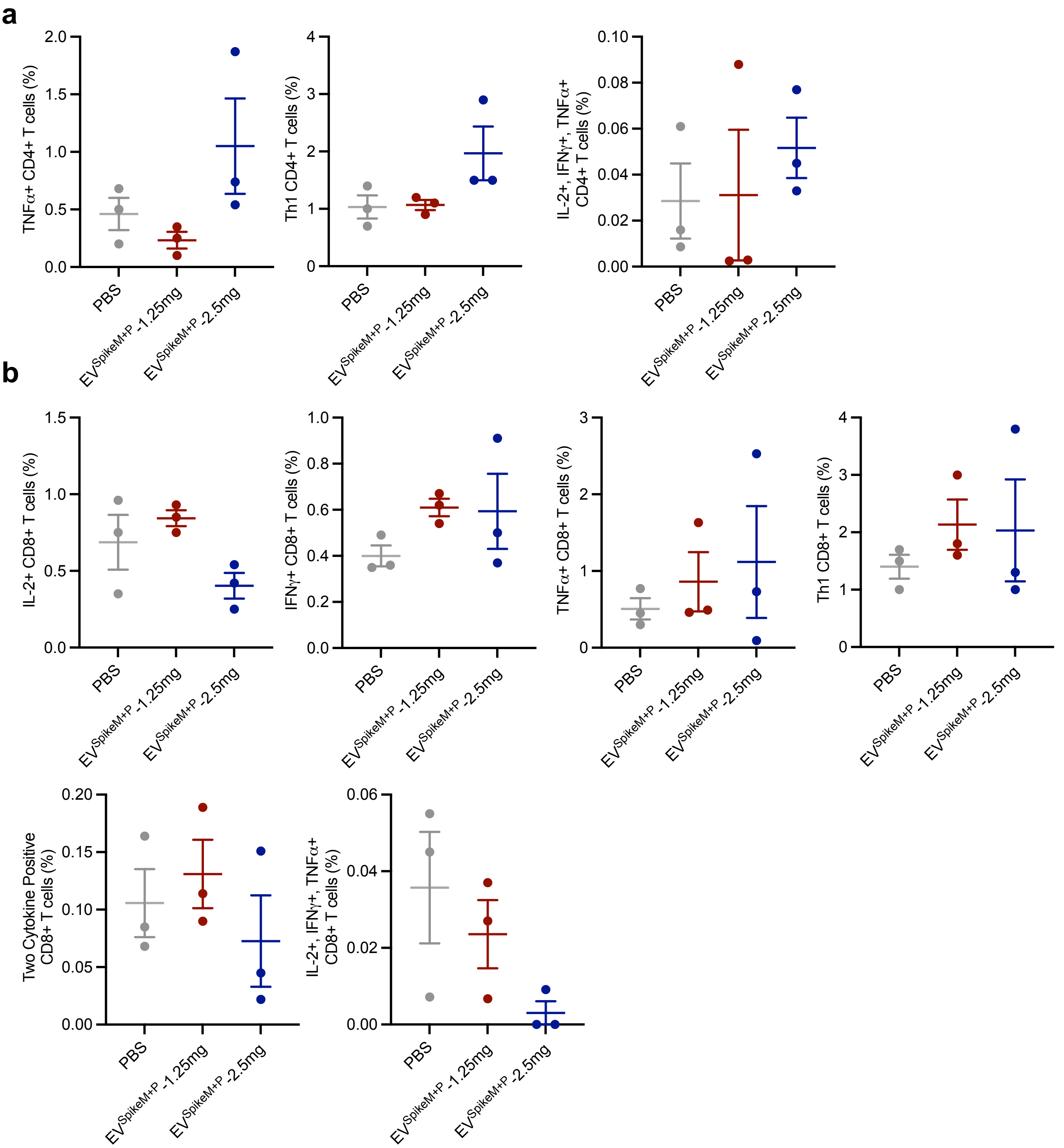

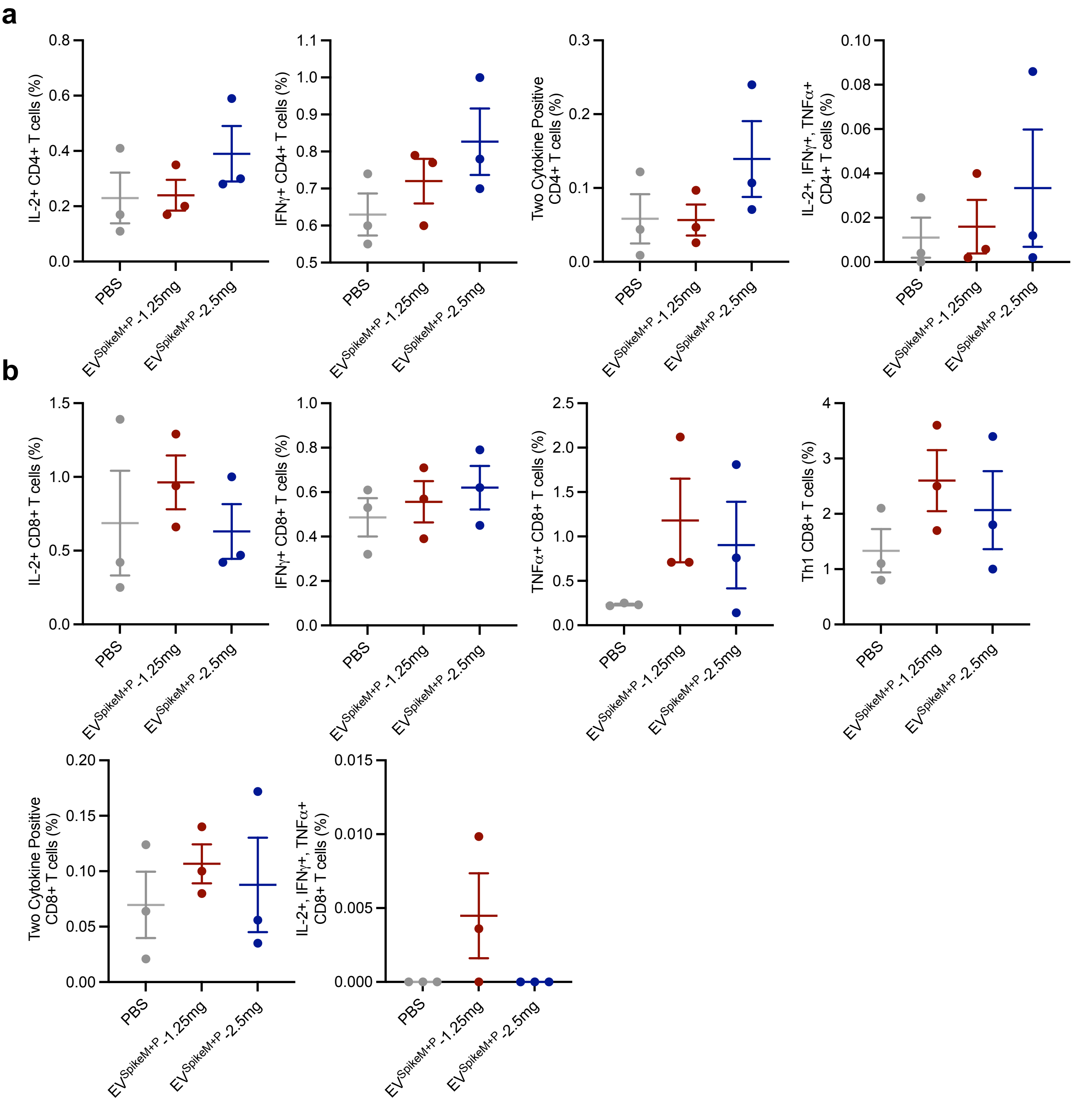
